# Multi-omics comparative analyses of synucleinopathy models reveal distinct targets and relevance for drug development

**DOI:** 10.1101/2025.06.20.660336

**Authors:** Britney N Lizama, Rick Shin, Hilary A North, Gary Look, Aidan Reaver, Kiran Pandey, Duc Duong, Nicholas T Seyfried, Anthony O Caggiano, Mary E Hamby

**Affiliations:** Cognition Therapeutics, Pittsburgh, Pennsylvania, USA; EmTheraPro Inc, Systems Biology, Atlanta, Georgia, USA; Emory University School of Medicine, Biochemistry, Atlanta, Georgia, USA

**Keywords:** Drug discovery, Preclinical models, Proteomics, Therapeutics

## Abstract

**Background:** The discovery and development of therapeutics for Parkinson’s disease (PD) requires preclinical models and an understanding of the disease mechanisms reflected in each model is crucial to success.

**Objective:** To illuminate disease mechanisms and translational value of two commonly utilized rat models of synucleinopathy – AAV-delivered human mutant hA53T alpha synuclein (α-Syn) and α-Syn preformed fibril (PFF) injection – using a top-down, unbiased, large-scale approach.

**Methods:** Tandem mass tag mass spectrometry (TMT-MS), RNA sequencing, and bioinformatic analyses were used to assess proteins, genes, and pathways disrupted in rat striatum and substantia nigra. Comparative analyses were performed with PD drug candidate targets and an existing human PD and dementia with Lewy body (DLB) proteomics dataset.

**Results:** Unbiased proteomics identified 388 proteins significantly altered by hA53T-α-Syn and 1550 by PFF-α-Syn compared to sham controls. Pathway and correlation analyses of these revealed common and distinct pathophysiological processes altered in each model: dopaminergic signaling/metabolism, mitochondria and energy metabolism, and motor processes were disrupted in AAV-hA53T-α-Syn, while immune response, intracellular/secretory vesicles, synaptic vesicles, and autophagy were more impacted by PFF-α-Syn. Synapses, neural growth and remodeling, and protein localization were prominently represented in both models. Analyses revealed potential biomarkers of disease processes and proteins and pathways also altered in patients, elucidating drug targets/ disease mechanisms the models best reflect.

**Conclusions:** Alignment of unbiased multi-omics analyses of AAV-hA53T and PFF-α-Syn models of synucleinopathy with PD and DLB patient data and PD drug development pipeline candidates identifies optimal models for testing novel therapeutics based on biological mechanisms.

## INTRODUCTION

Parkinson’s disease (PD) is characterized by selective degeneration of dopaminergic neurons in the substantia nigra pars compacta and a corresponding loss of nigrostriatal dopaminergic terminals.^1^ Accumulation of α-synuclein (α-Syn) in the form of Lewy bodies is the most prominent pathological feature of the disease,^1,2^ with phosphorylation of α-Syn at Ser129 (pS129-α-Syn) inducing cellular damage through aggregation, neurotoxicity, and extensive propagation.^3^

Many preclinical rodent models have been employed to investigate PD pathogenesis and potential therapies, including neurotoxin administration, genetic models, viral transfection of wildtype or mutant α-Syn, and intracerebral inoculation of α-Syn preformed fibrils/oligomers.^4^ Each approach exhibits altered cellular and molecular characteristics that recapitulate some aspects of human PD pathology. Two widely utilized approaches to studying the mechanisms that underlie synucleinopathy are the overexpression of human mutated A53T-α-Synuclein (hA53T-α-Syn) through adeno-associated virus (AAV)-mediated delivery to the striatum^5,6^ and the administration of preformed fibrils (PFF) of α-Syn (PFF-α-Syn) into the striatum or substantia nigra.^7,8^

As observed in an autosomal dominant form of PD, in which A53T-*SNCA* is genetically linked to familial PD, wherein patients express higher levels of mutant A53T-α-Syn,^9^ injection of AAV-A53T-α-Syn leads to overexpression of A53T-α-Syn transgene expression throughout the substantia nigra and striatum.^5^ The pathophysiological accumulation of α-Syn in nigral dopamine neurons as early as 4-6 weeks post-injection results in neuroinflammation and a loss of nigral dopaminergic neurons in the substantia nigra^10^ followed by production of toxic α-Syn oligomers and dystrophic neurites in the striatum, similar to Lewy neurite pathology in PD.^11^

In contrast, the intrastriatal α-Syn PFF model enables templating of endogenous non-mutant α-Syn that facilitates misfolding and aggregation of α-Syn monomers,^12–17^ recapitulating features of idiopathic PD including the presence of Lewy bodies, neuroinflammation in the striatum,^13,15^ and spread of the pathological pSer129-α-Syn^18^ and ensuing pathophysiological sequence of events over time to brain regions distal from the injection site.

Both preclinical models have facilitated understanding of the pathogenesis of synucleinopathies, particularly PD, and fostered the development of PD therapeutics. For instance, antisense oligonucleotides that target SNCA (Biogen)^19^ and leucine-rich repeat kinase 2 (LRRK2) (Ionis Pharmaceuticals)^20^ have been tested in PFF-α-Syn rodent models and shown to reduce production or levels of α-Syn and LRRK2, respectively. The biologic SAR446159 (Sanofi), an antibody under development for targeting α-Syn aggregates through immunotherapy, was shown to reduce α-Syn uptake by neurons and increase α-Syn clearance by microglia in preclinical PFF models.^21^ Another antibody, scFv, which targets a pathologic form of α-Syn, has been shown to improve motor deficits in the AAV-hA53T-α-Syn rodent model,^22^ as has the peptide GLP-1 analogue exendin-4.^23^ Small molecules including the mitochondrial division inhibitor-1 (mdivi-1)^24^ and the α-Syn aggregation inhibitor 20C^25^ have also been studied in hA53T-α-Syn preclinical models of synucleinopathy and shown to reduce neurodegeneration and improve motor deficits.

Despite the aforementioned successes in using these models, it is understood that there are many examples where tested therapeutics fail to demonstrate efficacy which is sometimes due to an incomplete understanding of the model’s utility for a given therapeutic mechanism. Thus, a better understanding of the disease mechanisms reflected in each model would improve efficiency for drug discovery programs testing therapeutics *in vivo*, and therefore accelerate delivery of medicines to patients.

To this end, we sought to holistically evaluate these two commonly used models through a large-scale discovery proteomics and transcriptomics approach. This unbiased, top-down approach complements the hypothesis-driven research on each of these models that has been carried out to date, producing a rich proteomics dataset that enabled the elucidation of proteins and pathways disrupted in each model. Comparative analyses with proteomes from postmortem brains of PD and dementia with Lewy body (DLB) patients^26^ delineated disease mechanisms (genes, proteins, and biological and molecular pathways) represented in each model, identifying their translational value. Further analyses were performed to dissect key mechanistic targets represented by each model, highlighting their strengths and limitations for use in preclinical drug discovery and development research. These findings serve as a resource or guide for model selection that may be valuable for investigators to increase the probability of success, expediting the advancement of therapeutics to the clinic.

## METHODS

### For detailed methods and procedures, see Supplemental Materials

#### Study design overview

Rats were used in one of two PD models: 1) AAV-mediated delivery of human A53T-α-Syn (or empty vector control) (Figure 1A, left) unilaterally into the SN, or 2) delivery of PFF-α-Syn (or monomeric α-Syn control) (Figure 1A, right) bilaterally into the striatum. At the end of the study (Day 43 (D43) for hA53T-α-Syn and EV controls, or D60 for PFF-α-Syn and monomeric α-Syn controls), brain tissue was collected and processed for postmortem analyses to validate model efficacy (Figure 1B) and for TMT-MS proteomics and RNA-seq (Figure 1C). For both models, animals were housed, fed, and acclimatized prior to procedures, and handled according to protocols approved by IACUC. Additional methods on the animal study design and procedures may be found in Suppl methods.

**Figure 1.**
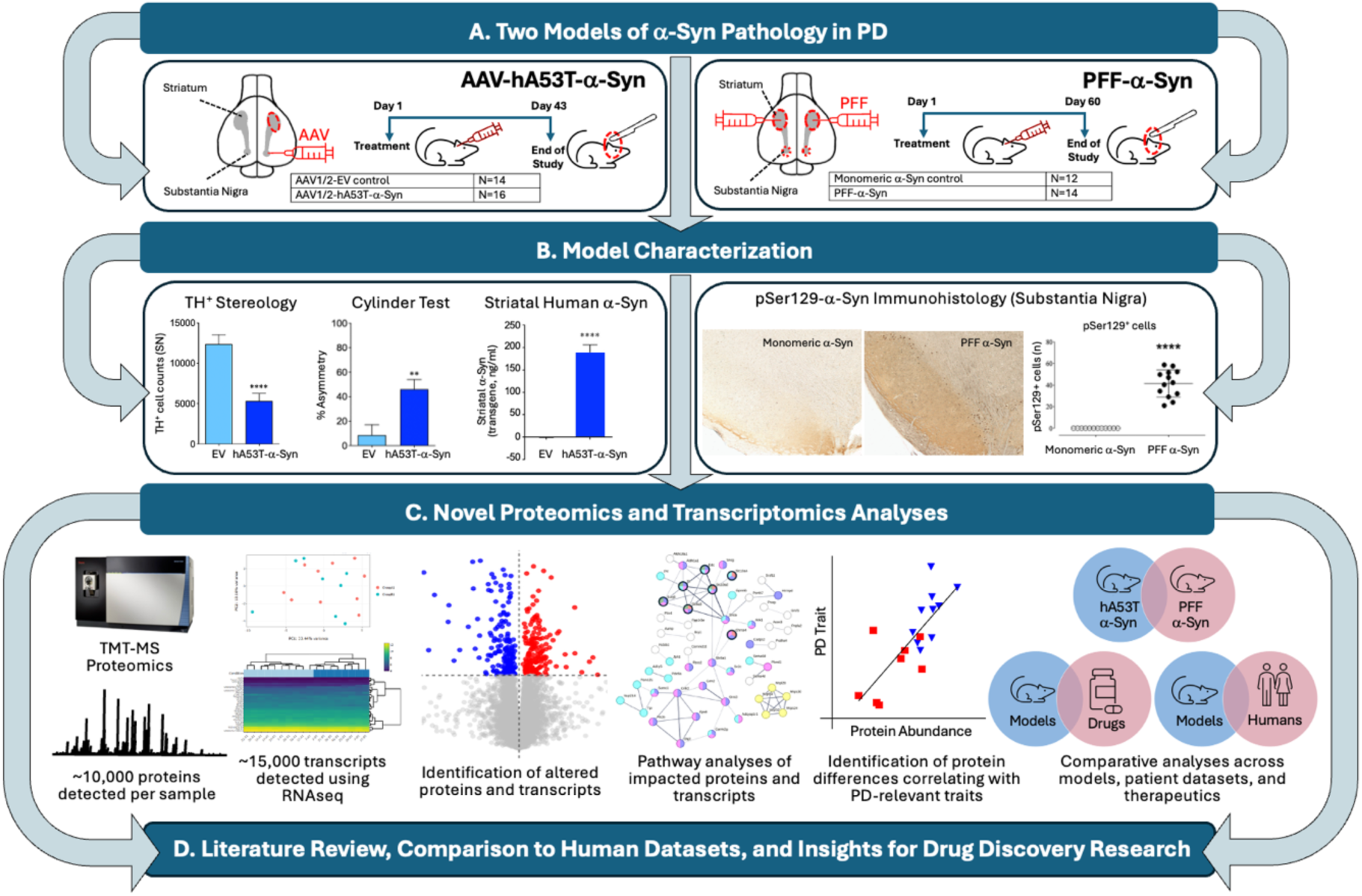
**Proteomic and transcriptomic phenotyping of two synucleinopathy models: AAV-hA53T-and PFF-**α**-Syn.** Parkinson’s disease (PD) was modeled in Sprague-Dawley rats and brain tissue was harvested for biochemical, histological and proteomic and transcriptomic analyses. **A. Left:** AAV-mediated stereotaxic delivery of human A53T α-Syn (hA53T-α-Syn): AAV-hA53T-α-Syn (n=16) or empty vector (EV) (n=14) was administered unilaterally into the rat substantia nigra (SN) on Day 1 (D1). Animals were sacrificed on D43 and tissue harvested. **Right:** Preformed fibril α-Syn (PFF-α-Syn) delivery: PFF-α-Syn (n=14) or monomer-α-Syn (n=12) was administered bilaterally into the rat striatum on D1. Animals were sacrificed on D60 and tissue harvested. **B. Left:** The impact of hA53T-α-Syn delivery was confirmed through nigral TH stereology, cylinder motor test, and striatal levels of human (transgene-derived) α-Syn via ELISA. **Right:** The impact of PFF-α-Syn delivery was confirmed through nigral α-Syn phosphorylated serine 129 (pSer129) immunohistochemistry. **C.** Proteomic and transcriptomic analysis workflow: Brain samples were processed and analyzed using tandem-mass tag mass spectrometry (TMT-MS) (striatum for both models) and RNA sequencing (striatum for AAV, SN for PFF). Differentially abundant proteins and RNA transcripts (hA53T- or PFF-α-Syn vs. controls, p≤0.05) were identified and assessed for functional enrichments, interconnectivity, and biological relevance using STRING, MetaCore, and Gene Ontology within and across models to assess similarities in differences between the two PD models, and correlation analyses were performed with classical PD hallmarks. **C-D.** Comparative analyses were performed to identify overlaps in a human PD and DLB patient proteomic dataset and overlaps with targets of PD and DLB therapeutics that are under development. Proteomic and transcriptomic insights gleaned herein can be used to facilitate model selection to guide researchers and foster drug development programs.

Differential abundances of proteins and genes were calculated and those achieving a change from baseline (p≤0.05) were analyzed using Gene Ontology (GO) to determine function, STRING and MetaCore for pathway analysis, comparative analysis to understand the differences between the two models, and correlation analyses to determine the proteins and genes associated with common PD-related traits (Figure 1C). Proteins, genes, and pathways determined to be impacted by hA53T-α-Syn and PFF-α-Syn delivery were considered in the context of known impacts of these models and therapeutic candidates for PD that are currently under development, to guide researchers in model selection for future studies (Figure 1C-D).

#### Brain tandem-mass tag mass spectrometry (TMT-MS) proteomics brief methods

All brain samples were balanced for treatment group across batches and loaded onto a 16-plex Tandem Mass Tag (TMTpro) kit (Catalog #A44520 and Lot #UI292951). Further, one Global Internal Standard (GIS) pool, created by aggregating lysate from all samples, was added per batch. Brain samples were processed and analyzed using Tandem-mass tag mass spectrometry (TMT-MS, see supplemental methods) followed by unbiased quantification. Using TMT-MS, 10274 proteins from the hA53T-α-Syn model cohort and 9802 proteins from the PFF-α-Syn model cohort were detected reliably across brain samples using the 50% detection exclusion criterion. One-way ANOVA followed by Tukey’s post hoc adjustment was performed on AAV-hA53T-α-Syn vs AAV-EV and on PFF-α-Syn vs monomer-α-Syn to identify differentially abundant proteins. Proteins with a p-value ≤0.05 were deemed significantly differentially abundant for each comparison.

#### RNA sequencing brief methods

Total RNA was extracted and high-quality samples were selected for sequencing analysis (see Supplemental Methods). RNAseq library preparation, sequencing, and bioinformatic analyses including differential expression analysis was performed at Azenta Life Science (South Plainfield, NJ, USA). Genes with a p-value ≤0.05 (Wald test) were deemed differentially expressed genes for each comparison.

#### Pathway and Comparative analyses of significantly impacted proteins and genes

##### Pathway analyses

Proteins and genes meeting the statistical threshold for each experiment (indicated for each figure) were assessed using STRING (version 11.5 or 12.0, indicated in each figure) and MetaCore™ (version 23.4.71500). Proteins without gene names (e.g., “0”) were excluded from STRING pathway analysis. The protein-protein interaction (PPI) network for each condition was exported and all pathway terms (e.g., for KEGG, GO Biological Processes, Reactome, etc.) were ranked according to strength, false discovery rate (FDR), or signal for interpretation, as indicated in each figure. For visualization of the PPI network, low, medium, and high confidence was used as appropriate (indicated for each figure). Unconnected nodes were excluded from the PPI maps.

##### GO terms categorization analysis

For Pie chart % representation, STRING Biological Process GO terms, Cellular Component GO terms, and Reactome terms were categorized according to discrete biologies and quantified accordingly (see Supplement).

##### Comparison to human PD and dementia with Lewy bodies (DLB) proteome dataset

We utilized previously published TMT-MS proteomics data from dorsolateral prefrontal cortex of PD or DLB patients (compared to age-matched healthy controls), derived from well-characterized autopsy collection (University of Pennsylvania Alzheimer’s Disease Research Center [ADRC]).^26^ Proteins meeting the statistical threshold of p≤0.05 for PD vs control or DLB vs control^26^ were considered for comparative analysis with significant differentially abundant proteins in the AAV-hA53T-α-Syn vs AAV-EV and in the PFF-α-Syn vs monomer-α- Syn analyses. Pathway analysis was performed using MetaCore™ (version 25.1.72000) and PPI maps were generated in STRING of the set of proteins overlapping.

## RESULTS

### Proteomic phenotyping of two mechanistically distinct rat models of synucleinopathy

#### AAV model

##### AAV-hA53T-α-Syn model validation

To ensure the integrity of the proteomic and transcriptomic phenotyping study, the classical PD hallmarks represented in the AAV-hA53T-α-Syn model – nigral dopaminergic neuronal loss, as assessed via tyrosine hydroxylase-positive (TH+) cell quantification, motor deficits as assessed via cylinder motor test, and elevated nigral α-Syn expression as assessed via ELISA – were measured 6 weeks following a unilateral injection of AAV-hA53T-α-Syn or AAV-empty vector (AAV-EV). (Figure 1A). TH^+^ cell counts were significantly decreased in the ipsilateral substantia nigra in AAV-hA53T-α-Syn injected compared to AAV-EV control rats (p<0.0001) (Figure 1B, left). Percent forelimb asymmetry, a measure of motor impairment given the unilateral injection, was also significantly elevated at 6 weeks in AAV-hA53T-α-Syn compared to AAV-EV controls (p<0.01). Levels of mutant human α-Syn expression were significantly elevated in the striatum compared to control (p<0.0001) (Figure 1B, left).

Additionally, striatal levels of DA transporter (DAT) (p<0.0001), DA (p<0.0001), DOPAC (p<0.0001) and the DA metabolite HVA (p<0.01) were also significantly lower (Supplemental Figure 1A-D), and striatal DA turnover (i.e., DOPAC + HVA / DA ratio) was significantly increased (p<0.05; Supplemental Figure 1E), in AAV-hA53T-α-Syn compared to AAV-EV control animals.

##### AAV-hA53T-α-Syn overexpression alters the striatal proteome and transcriptome

Nigral hA53T-α-Syn overexpression led to significantly altered expression levels of 388 proteins (202 decreased, 186 increased) relative to AAV-EV control (p≤0.05). Many identified proteins play key roles in dopamine synthesis and metabolism, neurotransmission, neural growth/ remodeling, and Parkinson’s disease including TH, vesicular monoamine transporter 2 (VMAT2; SLC18A2), DOPA decarboxylase (DDC), and neurexin 2 (NRXN2) (all decreased), as well as α-Syn (SCNA), apolipoprotein E (APOE), reticulon 4 receptor (RTN4R), and lysozyme 2 (LYZ2) (all increased) (Figure 2A, arrows; Table 1).

**Figure 2.**
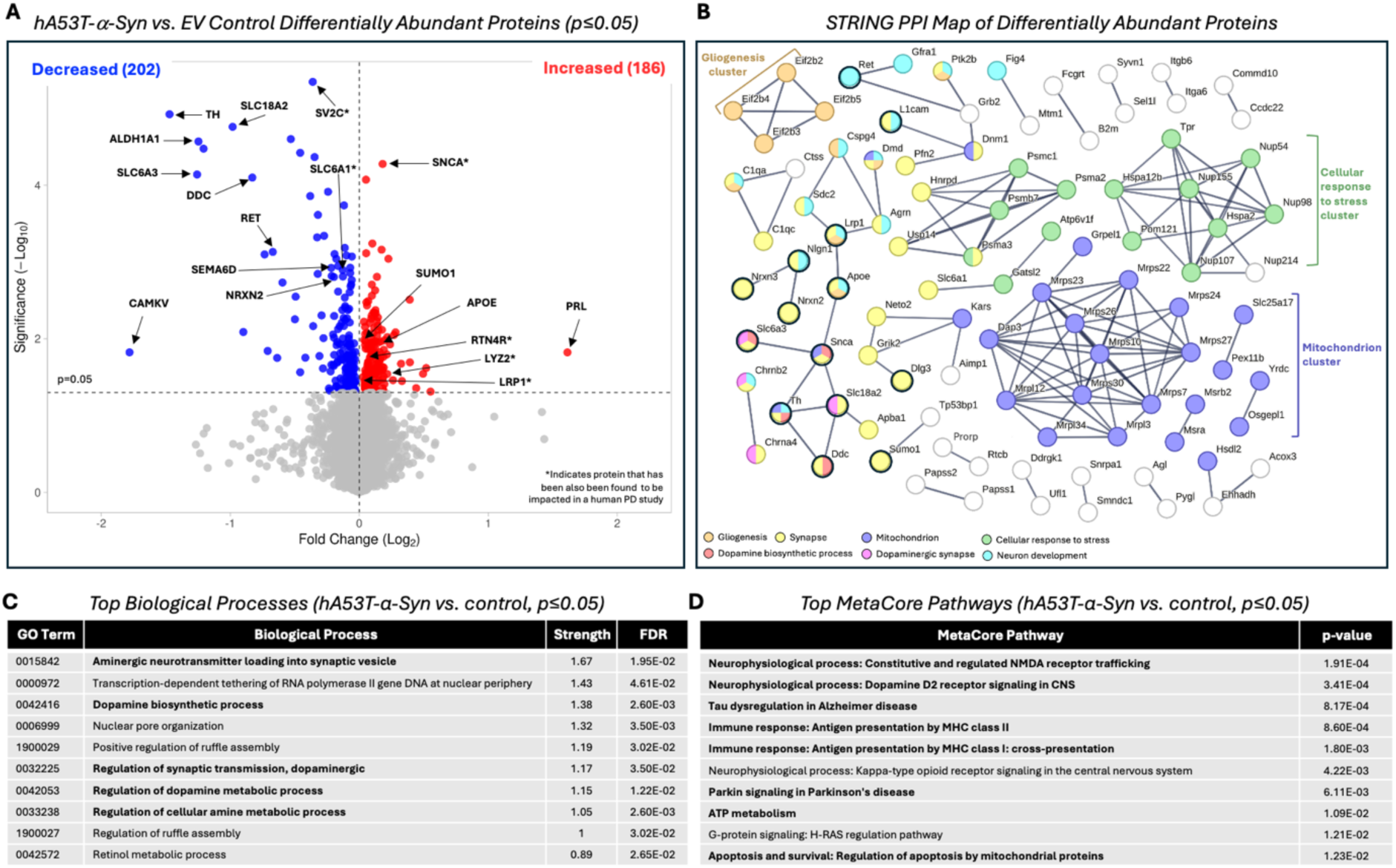
**AAV-hA53T-**α**-Syn expression alters striatal proteome and enriches pathways related to dopaminergic processes and synapses. A.** Differential abundance of striatal proteins measured through TMT-MS was assessed. Log2 fold change and significance for each protein detected can be visualized on the volcano plot. 388 proteins (202 decreased and 186 increased, relative to control), were found to be significantly (p≤0.05) differentially abundant. Proteins of interest are indicated with black arrows: ALDH1A1 (aldehyde dehydrogenase 1 family member A1), APOE (apolipoprotein E), CAMKV (calmodulin kinase-like vesicle-associated protein), DDC (dopa decarboxylase), SEMA6D (semaphorin 6D), SV2C (synaptic vesicle glycoprotein 2c), TH (tyrosine hydroxylase), LRP1 (LDL receptor related protein 1), PRL (prolactin), SLC18A2 (solute carrier family 18 member A2), SLC6A3 (solute carrier family 6 member 3), SLC6A1 (solute carrier family 6 member 1), SNCA (alpha-synuclein), SUMO1 (small ubiquitin like modifier 1), RET (ret proto-oncogene), NRXN2 (neurexin 2), and RTN4R (reticulon 4 receptor). *Indicates select protein that has been also been found to be impacted in the UPENN human PD cohort (Shantaraman et al., 2024): SV2C, SNCA, SLC6A1, LRP1, LYZ, RTN4R. B. STRING (v.11.5) Protein-Protein Interaction (PPI) map (highest confidence, 0.900; disconnected nodes hidden) of significantly (p≤0.05) differentially abundant proteins (hA53T-α-Syn vs. EV control). Clusters of proteins related to functionally enriched pathways and processes are indicated by color; proteins of interest circled in black. Neuron development, Gliogenesis, Dopamine biosynthetic process are Biological processes; Synapse, Mitochondrion, Dopamine synapse are Cellular Component; Cellular response to stress is Reactome. **C-D.** Pathway analyses using both STRING (GO Biological Processes) (**C**) and MetaCore mapping (**D**) show the processes and pathways most significantly disrupted in hA53T-α-Syn vs. EV control rats.

**Table 1:**
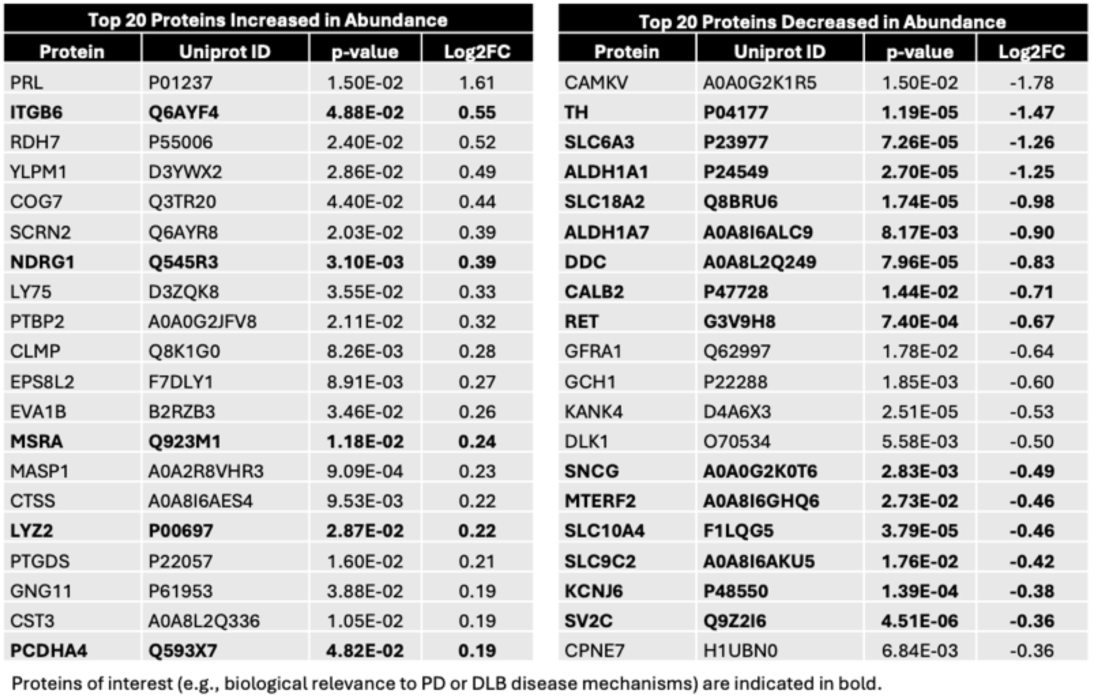
Top increased and decreased proteins in AAV-hA53T-α-Syn vs. EV control.

A STRING Protein-Protein Interaction (PPI) map (Figure 2B) shows interconnectivity among the differentially abundant proteins disrupted in the AAV-hA53T-α-Syn model, with prominent functional clusters in neurogenesis and gliogenesis, stress response, synapses – particularly dopaminergic synapses and dopamine synthesis – and mitochondria. Unbiased pathway analysis identified that the top ranked STRING Biological Processes (Figure 2C), via gene ontology (GO) analysis, as well as the MetaCore predesignated pathways (Figure 2D) disrupted were dopaminergic synaptic transmission, ATP/ mitochondrial processes, and immune response.

The impact of AAV-hA53T-α-Syn overexpression on the striatal transcriptome was investigated using RNAseq. Unbiased pathway analysis of the 611 significantly differentially expressed transcripts revealed the top ranked STRING Biological Processes (Supplemental Figure 2A) and top ranked MetaCore pathways (Supplemental Figure 2B) disrupted were synaptic activity and reorganization, and other cell signaling pathways. A comparative analysis of the 388 significantly impacted proteins and the 611 significantly impacted transcripts revealed neurodegeneration, apoptosis and cell survival, and neurotransmission to be represented biologies of the 28 overlapping transcripts and proteins affected in this model (Supplemental Figure 2C, D).

Proteomic correlates of TH^+^ cell counts in hA53T-α-Syn rat brain Given that TH^+^ cell numbers were significantly diminished in hA53T-α-Syn-expressing rat substantia nigra compared to EV controls (Figure 1B, left), and that the dopaminergic neurons of the substantia nigra send projections to the striatum, essential for proper motor function as part of the nigrostriatal pathway, a Pearson correlation analysis was performed to understand the relationship between substantia nigra TH^+^ cell counts and the altered striatal proteome (Figure 3). The most robustly correlated striatal proteins with substantia nigra TH^+^ cell counts were identified (Figure 3A), with some key disease-relevant proteins (i.e., related to dopamine synthesis, dopamine transport and PD etiology)^1,27–35^ described and plotted (Figure 3B, C). TMT-MS detected TH protein levels (-1.47 log2 fold change, p=1.19×10^-5^; Figure 2A) showed a robust positive correlation (r=0.73, p=6.40×10^-4^) with TH^+^ cell numbers in the substantia nigra (Figure 3A, B). The STRING PPI map illustrated interconnectivity of the most significant correlates (p≤0.01; 123 proteins), highlighting central hub clusters: synapses, including dopaminergic synapses, with widespread relevance to protein localization, neuron projection, and a mitochondrial cluster (Figure 3D). The top ranked STRING Biological Processes and Cellular Components relate overwhelmingly to synaptic and dopaminergic processes (Figure 3E).

**Figure 3:**
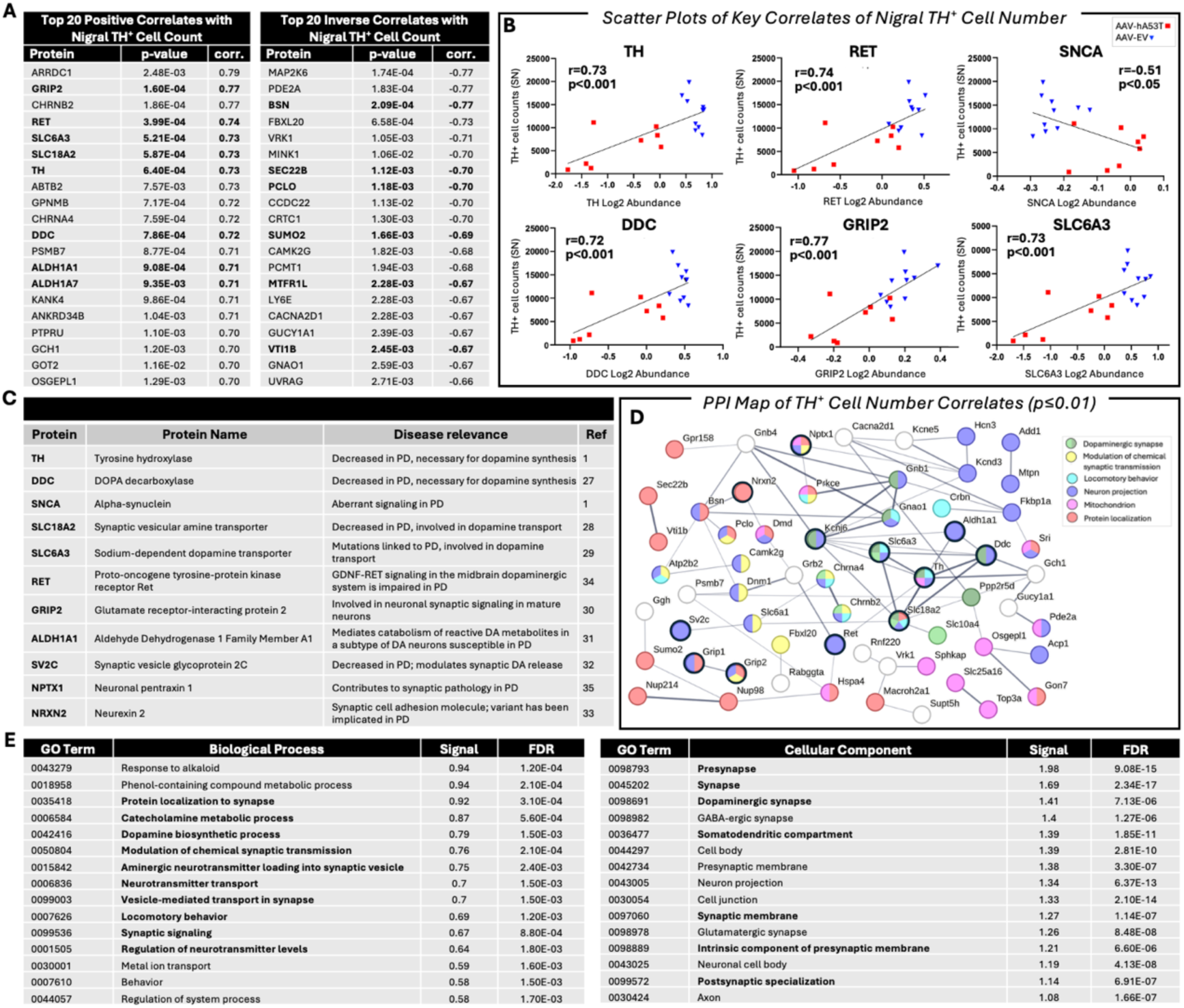
**Robust proteomic correlates of nigral tyrosine hydroxylase cell numbers identified in AAV- hA53T-**α**-Syn rat model. A.** Top 20 positively (left table) and inversely (right table) correlated proteins with tyrosine hydroxylase (TH)^+^ cell counts in the substantia nigra (SN) in hA53T-α-Syn-and EV control-treated rats. **B.** Scatter plots show correlation between TH^+^ cell count numbers and representative proteins in hA53T-α-Syn-(red) and EV control-treated (blue) rats. DDC: dopa decarboxylase; GRIP2: glutamate receptor interacting protein 2; RET: ret proto-oncogene; SLC6A3: solute carrier family 6 member 3; SNCA: synuclein alpha; TH: tyrosine hydroxylase. **C.** The known relationships of several key correlated proteins to Parkinson’s disease are described. **D.** Correlates (p≤0.01, positive and inverse) of nigral TH^+^ cell counts were analyzed using STRING (v.12.0) and the protein-protein interaction (PPI) map is shown (medium confidence 0.400, disconnected nodes hidden). TH, DDC, and SLC6A3 are central hubs in the PPI map, with prominent pathways relating to dopamine and synapses, locomotory behavior, neuron projection, mitochondrion, and protein localization indicated in color according to the legend. Proteins from **B** and **C** and others of biological or disease relevance are circled in black. **E.** Top (ranked by Signal) STRING Biological Process (left) and Cellular Component (right) gene ontology (GO) terms are shown.

#### PFF model

##### PFF-α-Syn rat model validation

To ensure proteomic and transcriptomic phenotyping were performed in a PFF-α-Syn model demonstrating the PD hallmark classically represented in this model,^13^ phosphorylated α-Syn at serine 129 (pSer129-α-Syn) was assessed via immunohistochemistry in substantia nigra brain sections and pSer129-α-Syn-positive (pSer129-α-Syn^+^) cells were quantified 60 days post-PFF-α-Syn or control injection. Indeed, the number of nigral pSer129-α-Syn^+^ cells was significantly elevated in PFF-α-Syn rats compared to monomeric α-Syn control (p<0.0001) (Figure 1B, right). In the parietal and insular cortices, pSer129-α-Syn immunohistological distribution and intensity levels were significantly elevated (p<0.0001) (Supplemental Figure 1G, H). Striatal dopamine was significantly reduced (Supplemental Figure 1F), as was DAT (Figure 7A); but DOPAC, HVA, and dopamine turnover were unchanged (data not shown).

##### PFF-α-Syn injection alters striatal proteome and nigral transcriptome

Post-intra-striatal PFF-α-Syn injection, a total of 1550 proteins were differentially expressed (775 decreased and 775 increased relative to monomeric control) (p≤0.05) (Figure 4A). A number of proteins relevant to neurotransmission, synaptic processes, neuroinflammation, and PD biology were significantly increased or decreased, including vesicle-associated membrane protein-2 (VAMP2, down), synaptotagmin-1 (SYT1, down), neuronal pentraxin-1 (NPTX1, down), gliomedin (GLDN, down), excitatory amino acid transporter 2 (SLC1A2, down), the protease HTRA1 (up), glial fibrillary acidic protein (GFAP; up), S100 calcium binding protein B (S100B, up), (Figure 4A, arrows; Table 2). The STRING PPI map (Figure 4B) demonstrated interconnectivity among altered proteins, with prominent functional clusters relating to autophagy, vesicle-mediated transport, synaptic vesicles, and the proteasome. The top ranked STRING Biological Processes (Figure 4C) and top ranked MetaCore pathways (Figure 4D) show that synapses/ synaptic vesicles and neuronal remodeling-related processes are predominant. The impact of PFF-α-Syn overexpression on the substantia nigra transcriptome was also investigated using RNAseq (Supplemental Figure 2). Unbiased pathway analysis of the 562 significantly (p≤0.05) differentially expressed transcripts revealed the top ranked STRING Biological Processes (Supplemental Figure 2E) and top ranked MetaCore pathways (Supplemental Figure 2F) disrupted were related to Notch signaling and immune response.

**Figure 4:**
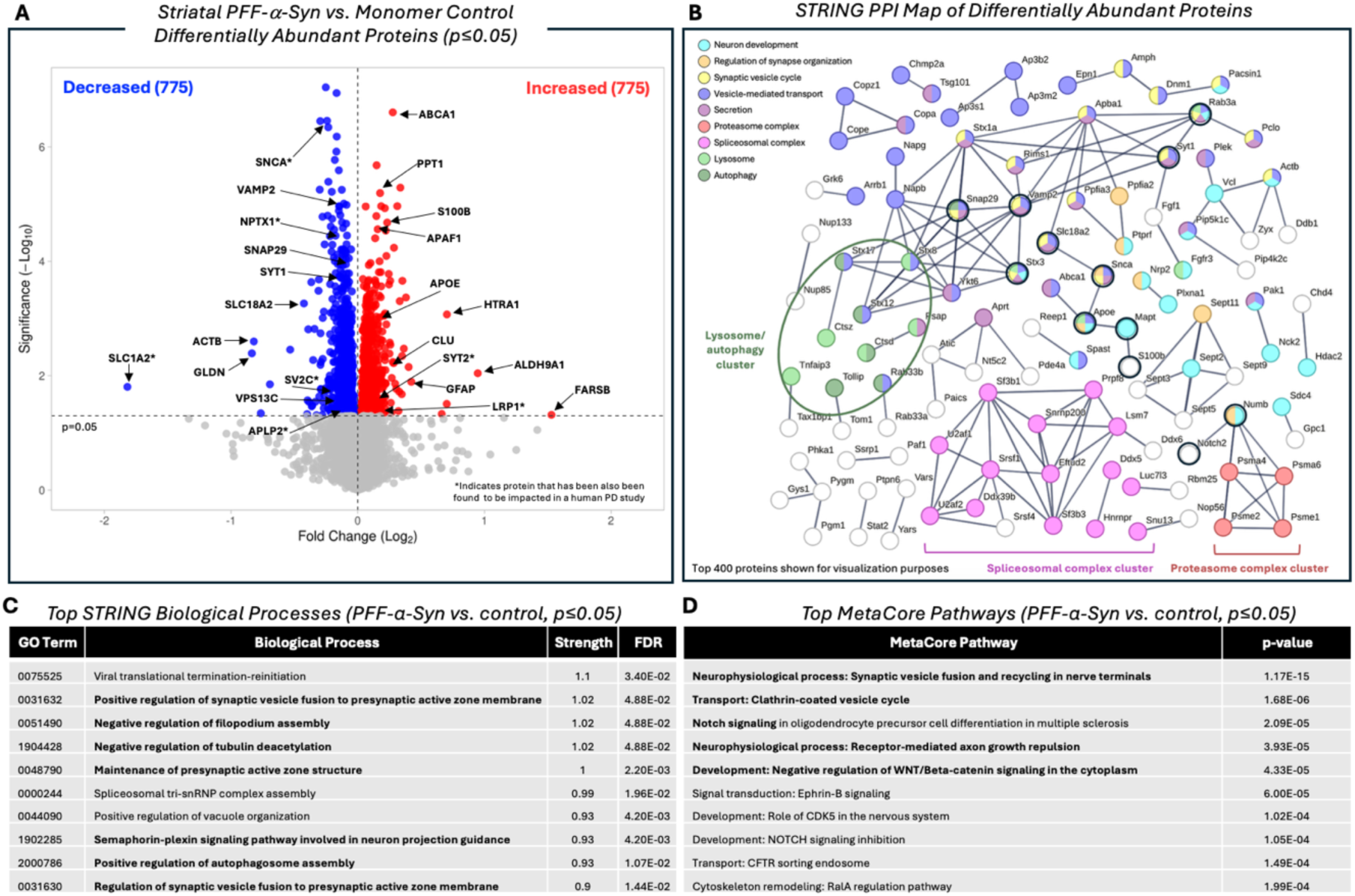
**PFF-**α**-Syn alters striatal proteome and impacts pathways related to vesicular trafficking, cell signaling, and axonal growth. A.** Differential abundance of striatal proteins measured through TMT-MS was assessed via ANOVA (PFF-α-Syn vs. monomer-α-Syn control; p≤0.05). Log2 abundance and significance for each protein detected is shown on the volcano plot. 1550 proteins (775 decreased and 775 increased relative to control), were found to be significantly (p≤0.05) differentially abundant. Proteins of interest are indicated with arrows: ABCA1 (ATP binding cassette subfamily A member 1), ACTB (beta actin), ALDH9A1 (aldehyde dehydrogenase 9 family member A1), APAF1 (apoptotic peptidase activating factor 1), APLP2 (amyloid beta precursor like protein 2), APOE (apolipoprotein E), CLU (clusterin), FARSB (phenylalanyl-tRNA synthetase subunit beta), GFAP (glial fibrillary acidic protein), GLDN (gliomedin), HTRA1 (HtrA serine peptidase 1), LRP1 (LDL receptor related protein 1), NPTX1 (neuronal pentraxin 1), PPT1 (palmitoyl-protein thioesterase 1), S100B (S100 calcium binding protein B), SLC1A2 (solute carrier family 1 member 2), SLC18A2 (solute carrier family 18 member A2), SNAP29 (synaptosome associated protein 29), SNCA (alpha synuclein), SV2C (synaptic vesicle glycoprotein 2c), SYT1 (synaptotagmin 1), SYT2 (synaptotagmin 2), and VAMP2 (vesicle-associated membrane protein 2). *Indicates select protein that has also been found to be impacted in a human PD study: APLP2, LRP1, NPTX1, SLC1A2, SNCA, SV2C, SYT2 (Shantaraman et al., 2024). **B.** STRING (v.11.5) Protein-Protein Interaction (PPI) map (top 400 most significantly altered proteins included for visualization purposes; highest confidence, 0.900; disconnected nodes hidden) of significantly (p≤0.05) differentially abundant proteins (PFF-α-Syn vs. monomeric control). Clusters of proteins related to functionally enriched pathways and processes are indicated by color and brackets. Proteins of interest are circled in black (MAPT, microtubule associated protein tau; NOTCH2, notch receptor 2; NUMB, NUMB endocytic adaptor protein; RAB3A, RAB3A, member RAS oncogene family; STX3, syntaxin 3). Synaptic vesicle cycle, Secretion, Regulation of synapse organization, Autophagy, Vesicle mediated transport, and Neuron development are Biological processes. Lysosome and spliceosome are Cellular component. Proteasome is a subcellular localization/ COMPARTMENT. **C-D.** Pathway analyses using both STRING **(C)** and MetaCore **(D)** mapping show the pathways most significantly altered in PFF-α-Syn vs. monomeric control rats. Pathways bolded show abundance of pathways related to cell signaling, synaptic/vesicular trafficking, and axonal growth.

**Table 2:**
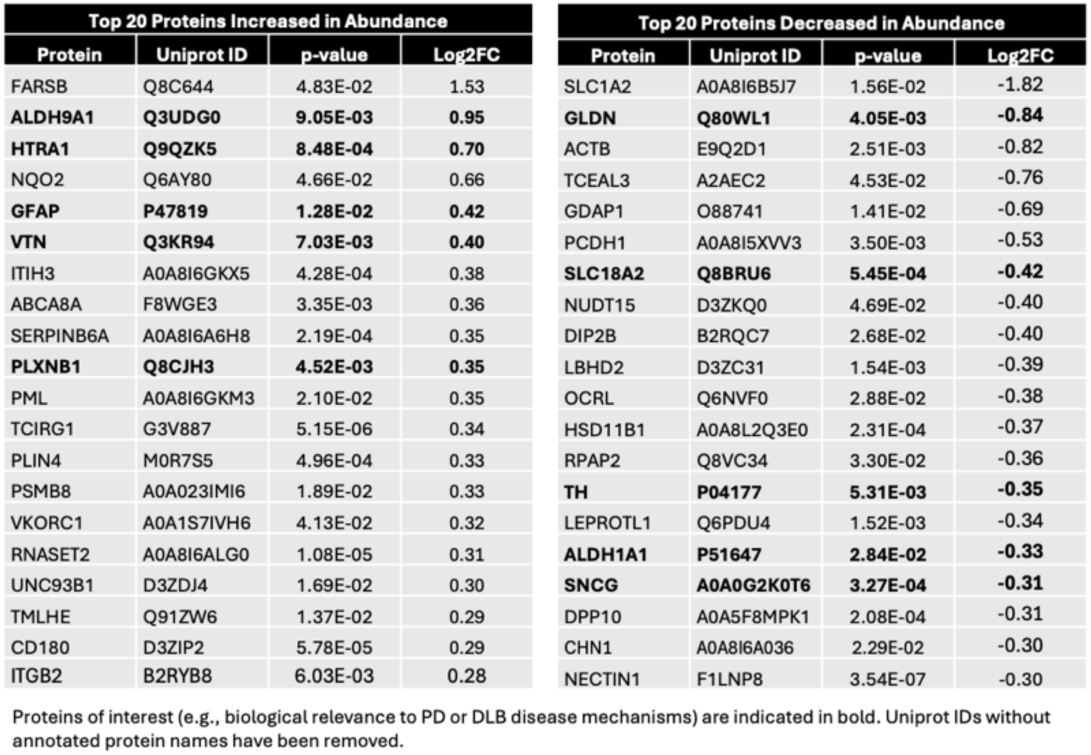
Top increased and decreased proteins in PFF-. α**-Syn vs. monomer control**

Proteomic correlates of pSer129-α-Syn^+^ cell counts in PFF-α-Syn rat brain Because hyperphosphorylation of α-Syn at serine 129 (pSer129-α-Syn) is a hallmark of synucleinopathies including PD and a key endpoint in the PFF model, we identified proteins robustly correlated with pSer129-α-Syn^+^ cell counts in SN as assessed via immunohistochemistry (Figure 5A). The 447 proteins that were found to be significantly (p≤0.01) correlated with pSer129-α-Syn^+^ cell counts were analyzed using STRING (Figure 5B, C). Vesicle-associated membrane protein 2 (VAMP2) and members of the vesicle/SNARE-related syntaxin family are central hubs of the PPI map (Figure 5B, green circle). The top ranked Biological Process and Cellular Component GO terms were overwhelmingly synaptic in nature: synaptic vesicles/ transport, secretion/ exocytosis, neurotransmission (Figure 5C).

**Figure 5:**
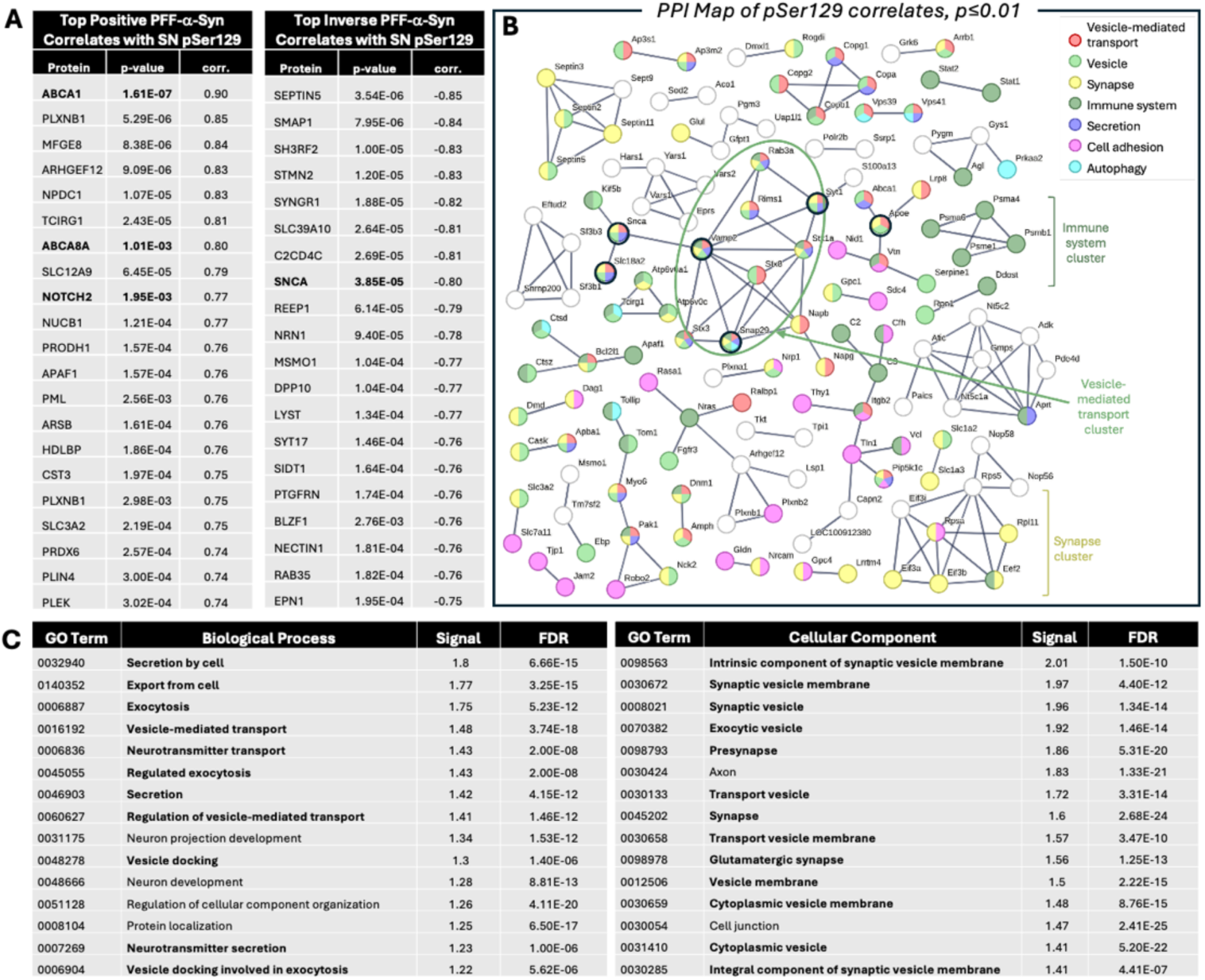
**Analysis of proteins correlated with pSer129-**α**-Syn in the PFF-**α**-Syn model. A.** Top 20 positively (left table) and inversely (right table) correlated proteins with phosphorylated serine 129 α-Syn (pSer129) levels in the substantia nigra in monomeric-and PFF-α-Syn rats. **B.** Correlates (p≤0.01, positive and inverse correlations) were analyzed using STRING (v.12.0) and the PPI map is shown (highest confidence, 0.900; disconnected nodes hidden). Proteins of interest are circled in black. The post interconnected proteins, Hub proteins, can be viewed here. **C.** Top (ranked by Signal) STRING Biological Process (left) and Cellular Component (right) gene ontology (GO) terms related to the correlates.

#### Across-model comparative analyses

##### Comparative analysis of proteins impacted in hA53T-and PFF-α-Syn models

To dissect the common molecular and functional phenotypes between the two models, an comparative analysis was performed of the 388 proteins altered in the AAV-hA53T-α-Syn model (vs. AAV-EV control, p≤0.05) and the 1550 proteins altered in the PFF-α-Syn model (vs. monomeric α-Syn control, p≤0.05), revealing 90 overlapping proteins disrupted in both models (Figure 6A). The top ranked MetaCore (Figure 6B) and STRING (Figure 6C, Biological Processes and Cellular Component GO terms) pathways indicate that both synucleinopathy models disrupt pathways involved in synapses and synaptic vesicles, including DA biosynthesis, locomotory behavior, neurogenesis, and transport. The STRING PPI map illustrates the connections between proteins and highlights chief functional clusters as: dopaminergic synapses, neurogenesis, transport, and locomotory behavior (Figure 6D).

**Figure 6:**
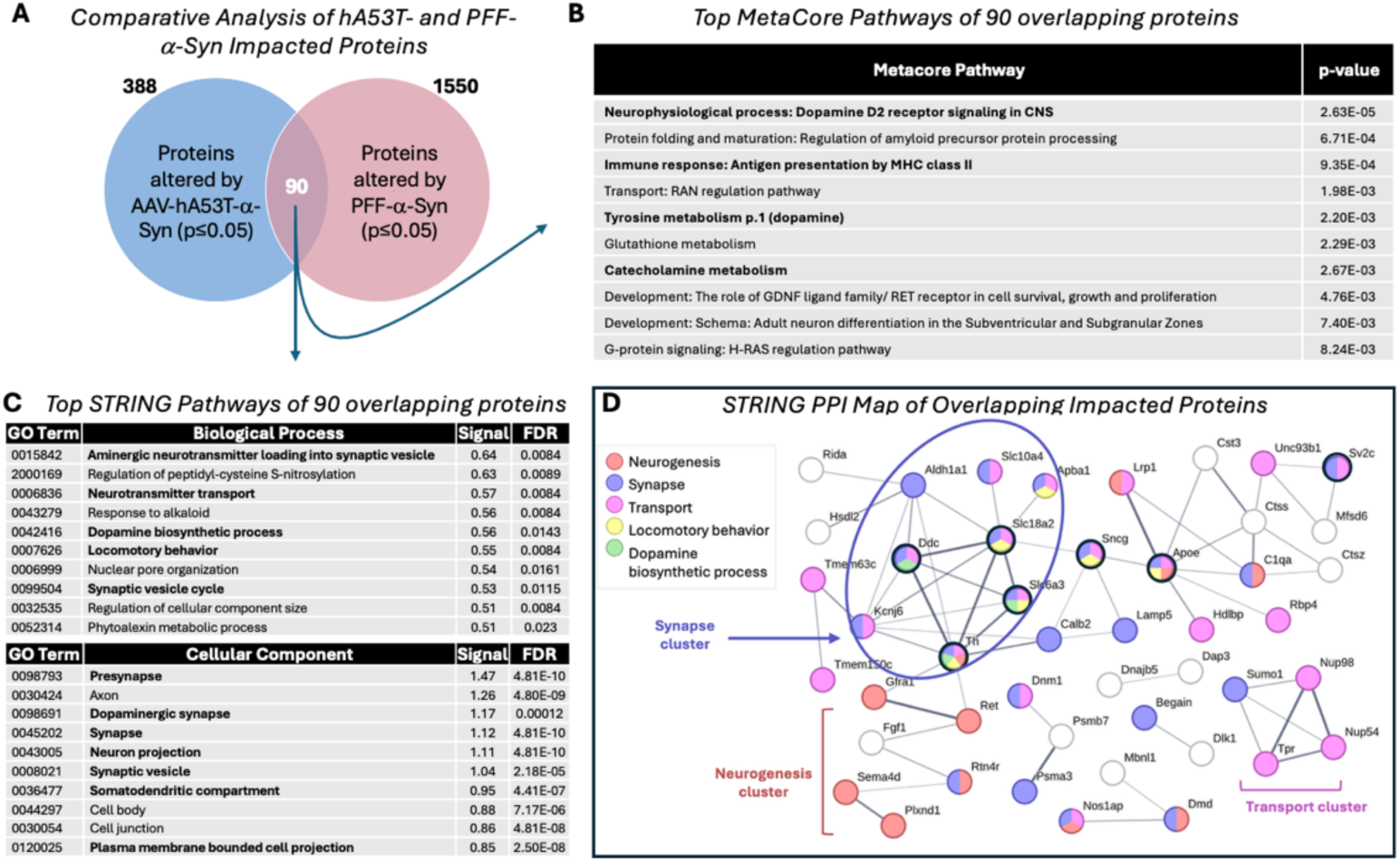
**Mechanistically similar disease biologies disrupted in both AAV-hA53T-and PFF-**α**-Syn rat models. A.** The proteins found to be significantly (p≤0.05) altered by AAV-hA53T-α-Syn and PFF-α-Syn (vs. respective controls) were comparatively analyzed to identify overlapping impacted proteins. The 388 proteins significantly altered by hA53T-α-Syn injection (blue circle, left) were found to have 90 overlaps (purple overlap, center) with the 1550 proteins significantly altered by PFF-α-Syn injection (red circle, right). **B-D.** Pathway analysis was performed on the 90 overlapping proteins. Disease-relevant pathways indicated in bold. **B.** The top 10 MetaCore pathways (ranked by FDR) found to be impacted. **C.** Using STRING (v.12.0) the top 10 (ranked by Signal) Biological Processes (top) and Cellular Components (bottom) found to be impacted. **D.** STRING protein-protein interaction map; medium confidence (0.400), disconnected nodes hidden. Proteins of interest are circled in black.

**Figure 7:**
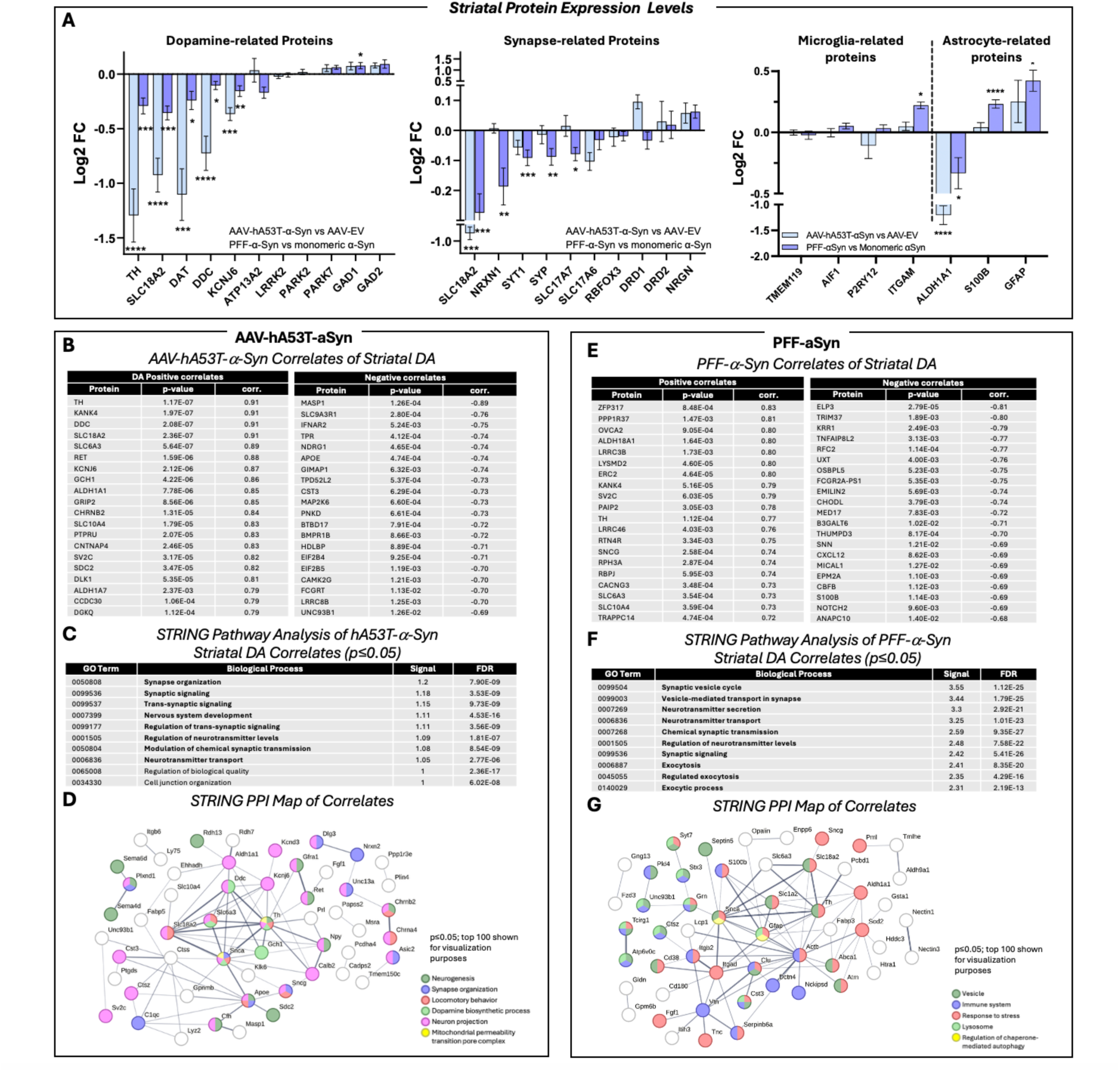
**Comparative analysis of proteins correlated with changes in dopaminergic indicators in AAV-hA53T-and PFF-**α**-Syn rat brain. A.** Dopaminergic neuronal markers (left), synapse-related proteins (middle), and microglia/ astrocyte-related markers (right) normalized to EV control (in AAV-hA53T-α-Syn-treated rats, light blue bars) or monomeric α-Syn (in PFF-α-Syn treated rats, dark blue bars), displayed in order from most decreased to most increased in abundance (log2 fold change, ±SEM). *Indicates significant. **B.** Top 20 positively (left) and inversely (right) correlated proteins with striatal DA levels in hA53T-α-Syn-and EV control-treated rats. **C-D.** Positive and inverse correlates (p≤0.05) of striatal dopamine were analyzed using STRING (v.12.0). Top (ranked by Signal) STRING Biological Process GO terms related to the correlates are shown (**C**). The protein-protein interaction (PPI) map **D**; top 100 proteins shown for visualization purposes, medium (0.400) confidence, disconnected nodes hidden). **E.** Top 20 positively (left) and inversely (right) correlated proteins with striatal dopamine levels in PFF-α-Syn-and monomer-α-Syn control-treated rats. Uniprot IDs without annotated protein names have been removed. **F-G.** Positive and inverse correlates (p≤0.05) were analyzed using STRING (v.12.0). Top (ranked by Signal) STRING Biological Process gene ontology (GO) terms related to the correlates are shown (**F**). The PPI map (**G**; top 100 proteins shown for visualization purposes, medium (0.400) confidence, disconnected nodes hidden).

##### Analysis of proteins correlated with dopamine levels in AAV-hA53T-and PFF-α-Syn rat brain

Overexpression of hA53T-α-Syn induces nigral dopaminergic cell death and thereby lowers dopamine levels in the striatum, whereas in the PFF-α-Syn model, a milder reduction in DA levels is observed.^12,36^ Consistent with prior reports, striatal dopamine levels were significantly decreased in hA53T-α-Syn overexpressing rats (Supplemental Figure 1B) and mildly, but significantly, reduced in PFF-α-Syn injected striatum (Supplemental Figure 1F) compared to control. Given the differences in pathology, we used proteomics in attempt to discern the biological underpinning of the dopamine deficiency induced by differing forms of aberrant α-Syn. To this end, a number of markers of dopaminergic cell number and function were assessed for both models (Figure 7A, left graph); and many, such as TH and DDC, essential for dopamine synthesis, were found to be significantly reduced in both models. These reductions could result from either a loss of striatal dopaminergic neurons themselves, of dopaminergic terminals from nigrostriatal projections, or from decreased dopamine synthesis. To probe these possibilities, protein levels of the neuronal cell body marker NeuN (RBFOX3) was evaluated, and no change was observed in either model compared to controls (Figure 7A, middle graph), suggesting the decrease was not due to a decrease in striatal neuron numbers. Subtle differences were found in synaptic marker protein levels: the PFF-α-Syn model had significantly decreased pre-synaptic proteins, which may be indicative of loss of synaptic terminals innervating the striatum, whereas post-synaptic markers levels were unchanged (Figure 7A, middle graph). In the hA53T-α-Syn model, most synaptic markers were unchanged except for the presynaptic monoamine neurotransmitter-specific VMAT2 (SLC18A2), which was significantly decreased (Figure 7A, middle graph).

Unbiased proteome-wide correlates with dopamine levels were identified and the top correlated proteins detected in AAV-hA53T-α-Syn overexpressing rats (and AAV-EV controls) are listed (Figure 7B). The 617 proteins significantly (p≤0.05) correlated with dopamine levels were analyzed using STRING (Figure 7C, D). TH, DDC, and dopamine transporter (SLC6A3 (DAT)) are central hubs in the PPI map, with synapse, neurogenesis, and dopamine strongly represented (Figure 7D). Similarly, the top ranked Biological Process GO terms reveal a strong emphasis on synapse and neural development (Figure 7C), with 7 out of 10 top affected Biological Processes being synapse-and neurotransmitter-related. Likewise, the top 20 positively and top 20 inversely correlated proteins, detected in PFF-α-Syn treated rats (and monomeric α-Syn controls), with striatal dopamine levels are listed (Figure 7E) and all 695 proteins that were found to be significantly (p≤0.05) correlated with dopamine levels were analyzed using STRING (Figure 7F, G). TH and GFAP are central hubs in the PPI map, along with S100b and clusterin (CLU), with vesicles, response to stress, and immune function strongly represented (Figure 7G). The top ranked Biological Process GO terms reveal a strong emphasis on synapse, vesicle, and secretion/ exocytosis (Figure 7F, 10 out of 10 top-affected Biological processes).

##### Comparative analysis of ‘pathways and processes’ impacted by hA53T-and PFF-α-Syn models

To further probe the biological processes and functional pathways impacted by either hA53T-or PFF-α-Syn or both models, the STRING-identified GO Biological Processes, GO Cellular Components, and Reactomes represented by each model were systematically grouped into functional categories (see methods) and quantification by percent of representation indicated. The Biological Process categories most impacted in hA53T-α-Syn (Figure 8, top row) included synapses and neurotransmission (22%) – including DA-specific synthesis and signaling (3%), as well as neural growth/ remodeling (19%), mitochondria/ metabolism (6%), and transport (16%)/ protein localization (9%). Synapses (20%) or vesicles (10%) and motor/ muscle (5%) were strongly represented GO Cellular Components of interest, and gene expression (20%) and mitochondria/ metabolism (15%) were amongst the top Reactome pathways impacted by the hA53T-α-Syn model (Figure 8, top row).

**Figure 8:**
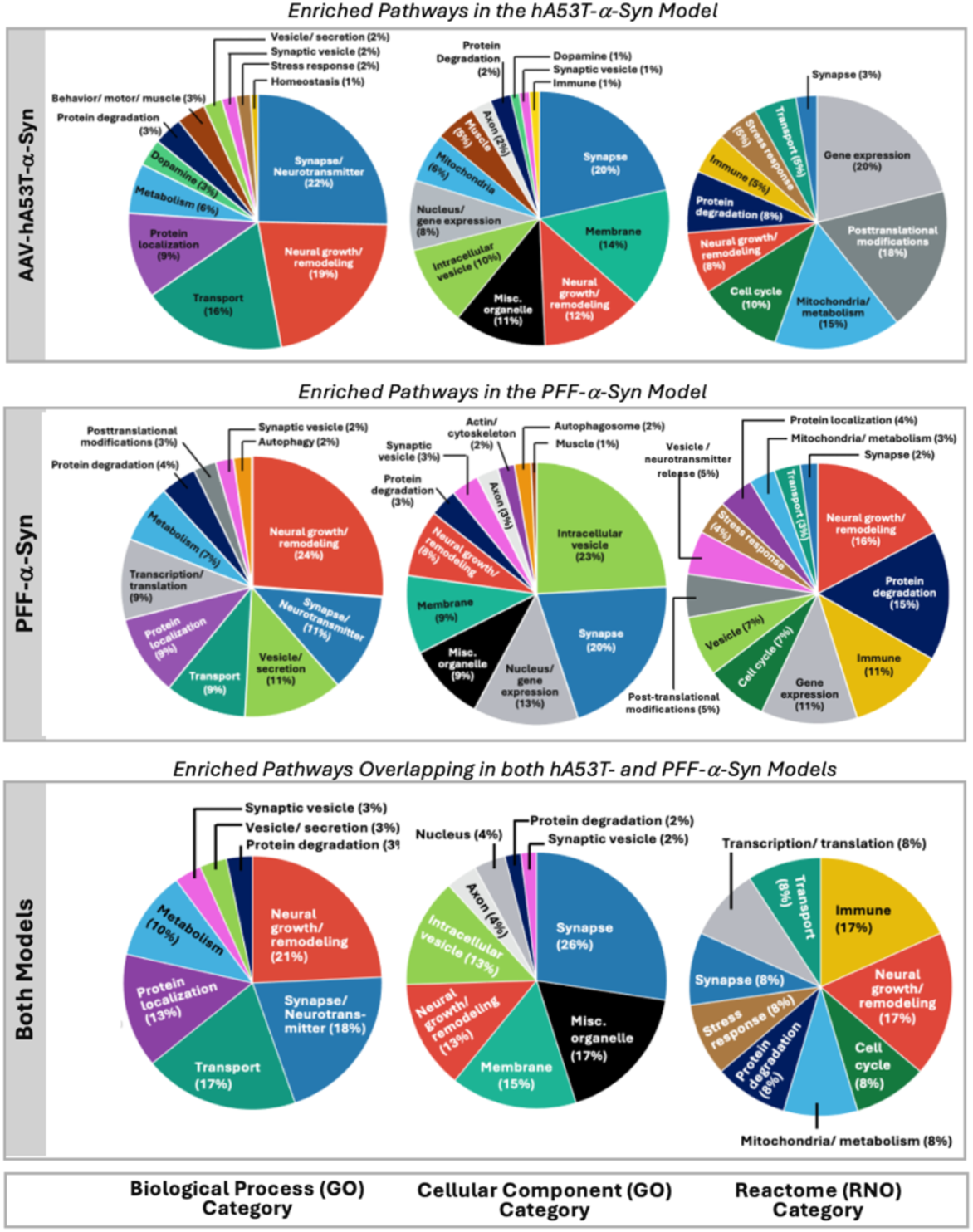
Analysis of enriched pathways and impacted processes in AAV-hA53T-and PFF-. α**-Syn rat brain proteomes.** STRING-identified pathways (Biological Process, Cellular Component, and Reactome terms, meeting a threshold of Signal≥0.3) impacted by hA53T-α-Syn (**Top row**) or by PFF-α-Syn (**Middle row**) were determined: functional categories including Synapse/ neurotransmission, Neural growth/ remodeling, Vesicle/ secretion, Protein degradation, Autophagy, etc. Quantifications of impacted pathways are shown for Biological Process (left column), Cellular Component (middle column), and Reactome (right column). **Bottom row:** Comparative analysis across models of enriched pathways and impacted processes: STRING-identified pathways (Biological Process, Cellular Component, and Reactome; Signal≥0.3) impacted by both hA53T-and PFF-α-Syn were analyzed comparatively to identify overlapping impacted pathways. The common pathways/ processes were assigned to categories as above, and % representation is shown.

For PFF-α-Syn (Figure 8, middle row), impacted GO Biological Process categories included neural growth/ remodeling (24%), synapse/ neurotransmission (11%), vesicle/ secretion (11%), and transport (9%) as well as protein localization (9%) were the top represented. GO Cellular Components more prominently impacted by PFF-α-Syn were intracellular vesicles (23%) and synapses (20%), and top Reactome pathways included neural growth and remodeling (16%), protein degradation (15%) and immune function (11%) (Figure 8, middle row).

Of 127 Biological Processes impacted in hA53T-α-Syn and the top 200 GO Biological Processes impacted in PFF-α-Syn, 70 were found to be overlapping. Of 84 Cellular Component GO terms impacted in hA53T-α-Syn and the top 150 Cellular Component GO terms impacted in PFF-α-Syn, 53 were found to be overlapping. Finally, of 40 Reactomes impacted in hA53T-α-Syn and the top 100 Reactomes impacted in PFF-α-Syn, 12 were found to be overlapping. The processes impacted in both models predominantly reflected neural growth/ remodeling (21%; Biological Process); synapses (26%; Cellular Component), and immune (17%, Reactome) function (Figure 8, bottom row).

#### Comparative analysis of hA53T-and PFF-α-Syn-impacted proteins and human patient proteome

##### Comparative analysis of AAV-hA53T-α-Syn and PFF-α-Syn models and human patient proteome

A comparative analysis of hA53T-αSyn with previously characterized postmortem human synucleinopathy tissue cohorts^26^ – PD and the second most prevalent synucleinopathy, dementia with Lewy body (DLB), was performed. Many of the significantly impacted proteins in the hA53T-and PFF-α-Syn striatum were disrupted in PD (Figure 9A, B) and DLB (Figure 9C, D) patient brain compared to age-matched healthy control brain. More specifically, a total of 46 significantly (p≤0.05) impacted proteins were common between the hA53T-α-Syn model and PD patient cohort (Figure 9A); MetaCore pathway analysis of these overlapping proteins revealed strong participation in neurite outgrowth, immune function, neurodegenerative pathological hallmarks, and protein folding related to neurodegenerative disorders. The STRING PPI map demonstrates the interconnectivity of the impacted proteins with SNCA, GABA transporter protein type-1 (SLC6A1), synaptic vesicle glycoprotein-2C (SV2C), and RTN4R among the most centrally connected nodes. Comparison of the hA53T-α-Syn impacted proteins with the DLB patient proteome from the same cohort revealed 73 common proteins (Figure 9C); MetaCore pathway analysis revealed immune and growth factor pathways to be strongly represented (Figure 9C). The STRING PPI map shows mitochondrial enzyme aldehyde dehydrogenase 18 family, member A1 (ALDH18A1), small ubiquitin-like modifier (SUMO1), and heat shock protein family A (HSP70) member-2 (HSPA2) to be centrally connected nodes among the overlapping proteins.

**Figure 9:**
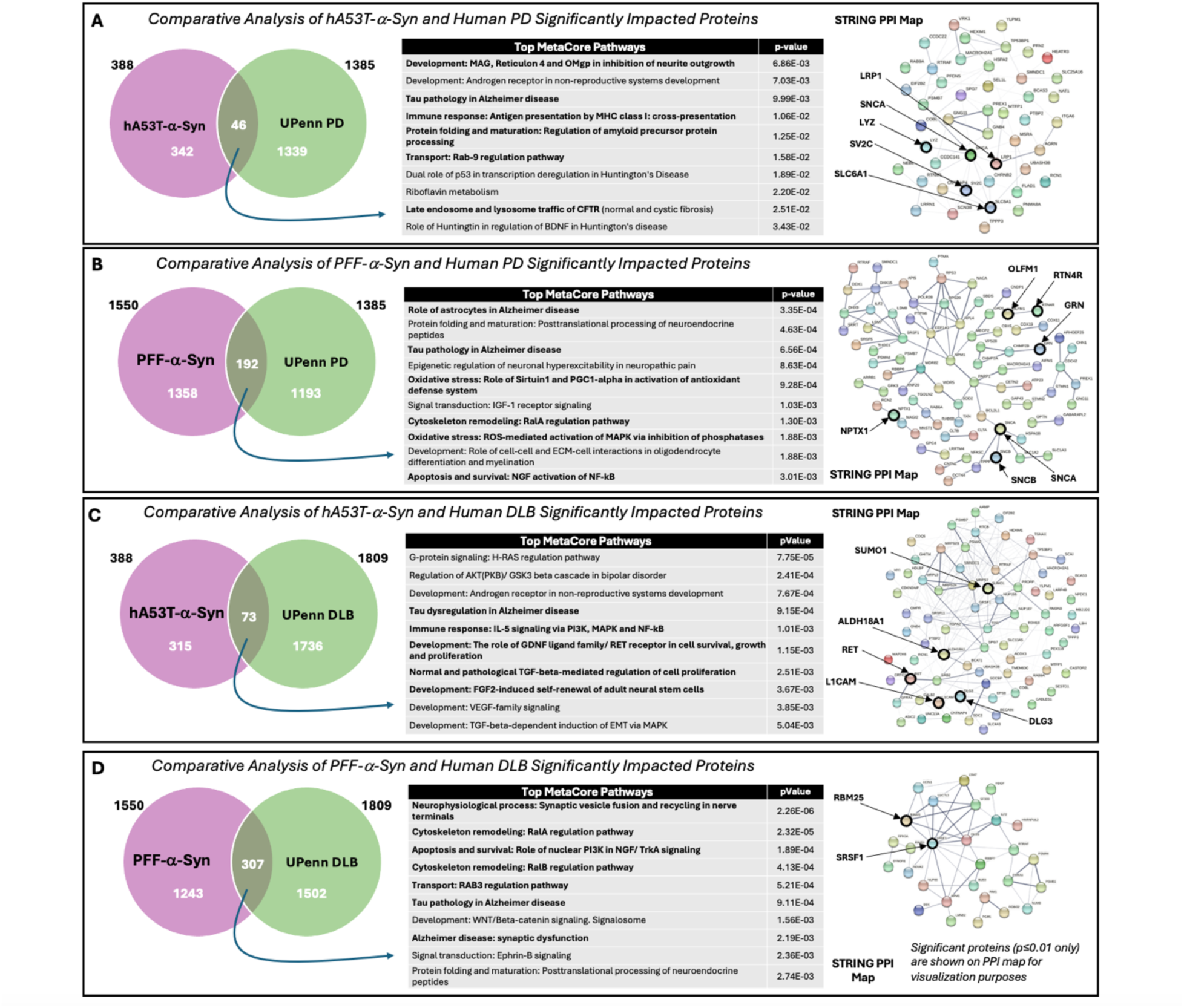
**Comparative analysis of hA53T-and PFF-**α**-Syn with human PD patient proteomes.** Comparative analysis of significantly impacted proteins in: **A**. the hA53T model (AAV-hA53T-α-Syn vs. EV control, p≤0.05, n=388) and in post-mortem brain from patients from the UPenn PD proteomic dataset (PD vs. healthy age-matched controls, p≤0.05, n=1385) revealed n=46 common impacted proteins (overlap of circles in Venn diagram), **B.** in the PFF model (PFF-α-Syn vs. monomer control, p≤0.05, n=1550) and human patients from the UPenn PD proteomic dataset (PD vs. healthy age-matched controls, p≤0.05, n=1385) revealed n=192 common impacted proteins (overlap of circles in Venn diagram), **C.** in the hA53T model (AAV-hA53T-α-Syn vs. EV control, p≤0.05, n=388) and in human patients from the UPenn DLB proteomic dataset (DLB vs. healthy age-matched controls, p≤0.05, n=1809) revealed n=73 common impacted proteins (overlap of circles in Venn diagram), **D.** in the PFF model (PFF-α-Syn vs. monomer control, p≤0.05, n=1550) and in human patients from the UPenn DLB proteomic dataset (DLB vs. healthy age-matched controls, p≤0.05, n=1809) revealed n=307 common impacted proteins (overlap of circles in Venn diagram). **A-D**. Top MetaCore pathways (table) and STRING PPI map (right; p<0.05, unless specified otherwise) are shown.

As with the hA53T-α-Syn significantly impacted proteins, the PFF-α-Syn significantly impacted proteins were compared to the PD and DLB postmortem patient brain tissue proteomes^26^ to identify commonly impacted proteins (Figure 9). A total of 192 significantly (p≤0.05) impacted proteins were found to be in common between the PFF-α-Syn model and PD patient cohort (Figure 9B). MetaCore pathway analysis of these overlaps revealed processes including astrocyte biology, oxidative stress, and apoptosis to be represented, and the STRING PPI map shows that SNCA, β-synuclein (SCNB), progranulin (GRN), nucleophosmin-1 (NPM1), and poly (ADP-ribose) polymerase-1 (PARP1) are centrally connected hubs. Individual overlapping impacted proteins of interest are indicated on the volcano plot in Figure 4A with an asterisk. Finally, comparison of the PFF-α-Syn impacted proteins with the DLB patient proteome from the same cohort revealed 307 common proteins (Figure 9D). MetaCore pathway analysis of these overlaps revealed processes including synapses, cytoskeleton remodeling, and apoptosis/ cell survival to be represented, and the STRING PPI map shows that Interleukin enhancer binding factor-2 (ILF2), histone-binding protein RBBP7, and serine/arginine-rich splicing factor-1 (SRSF1) are centrally connected hubs.

##### Comparative analysis with PD drug development pipeline targets

Understanding the disease mechanisms best reflected in each model is crucial to successful study design for PD drug development; thus, the multi-omics datasets were used to summarize the mechanisms convergent and distinct to each *in vivo* model. Results from the pathway analyses of the significantly impacted proteins, the correlation analyses for dopamine, tyrosine hydroxylase (hA53T-α-Syn) and pSer129 (PFF-α-Syn), the categorization analyses of impacted processes, and the comparative analyses with human patient proteomic data were qualitatively combined to determine the weighted representation of each disease-relevant process across models (Figure 10). Synapse and neurotransmission-related pathways, neural growth and remodeling, transport, protein localization, and stress response, were represented in the proteomes of both models (Figure 10, green). Pathways related to dopamine biology and mitochondrial function were more prominent in the hA53T-α-Syn model (Figure 10, orange), while pathways related to secretory vesicles and autophagy were distinctly represented in the PFF-α-Syn model (Figure 10, purple). To further refine these conclusions, the comparative analysis with the PD and DLB human patient cohorts was then considered in weighting the representation of each disease biology in the models (Figure 10, arrows indicating direction of weighting).

**Figure 10:**
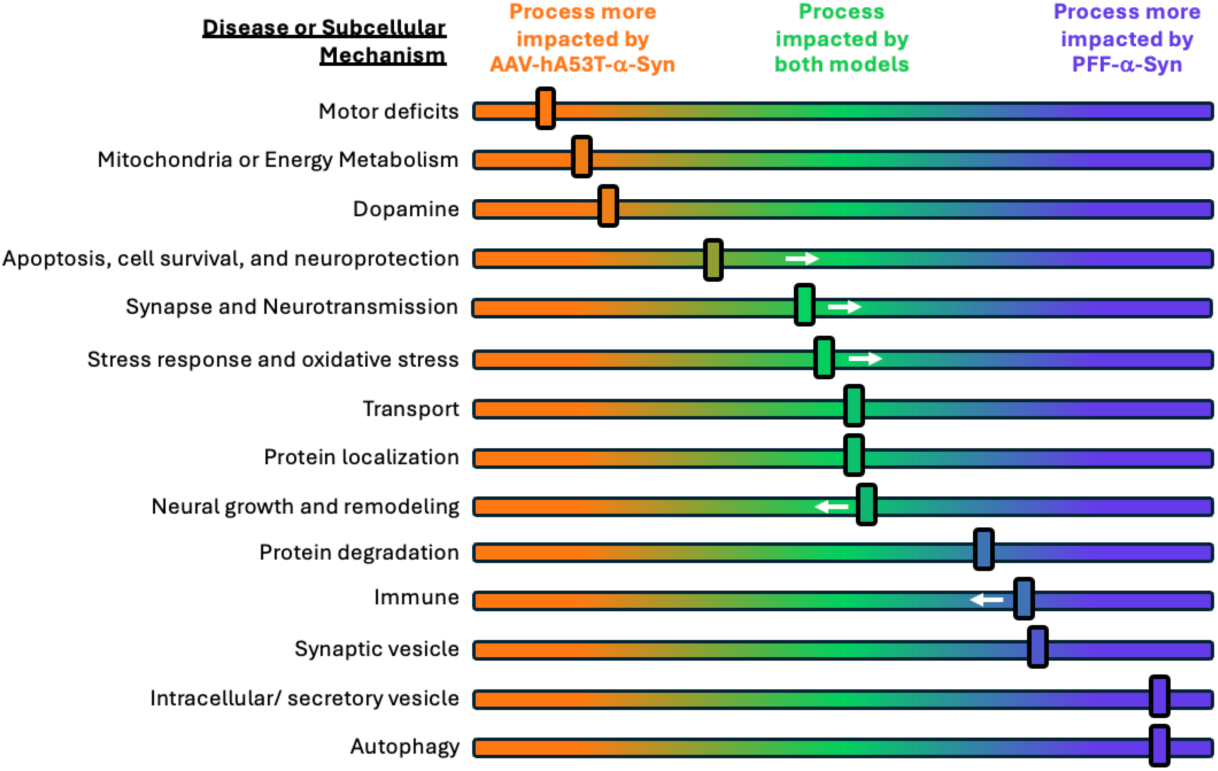
**Summary of disease-relevant processes impacted in AAV-hA53T-and PFF-**α**-Syn rat models of PD**. A number of biological, disease-related mechanisms were found to be impacted by the AAV-hA53T-and PFF-α-Syn rat models of synucleinopathy across the proteomic and transcriptomic analyses performed in this study. While many of these processes were represented to some degree in each model, some processes were more prominently impacted by hA53T-α-Syn (sliders weighted to orange) or by PFF-α-Syn sliders weighted to purple). Some processes appeared to be impacted fairly evenly by both models (sliders in middle/ green). White arrows indicate a weighting of process towards one model or the other based on human dataset overlap analysis in Figure 9.

To better understand the utility of each model in drug discovery, we asked whether hA53T-or PFF-α-Syn impacts levels of key targets of interest in current PD/DLB therapeutics under development.^37^ Log2 fold protein level changes in both models for a list of drug targets is shown (Figure 11A). For example, the endolysosome proteins GPNMB (decreased) and RILPL1 (increased) were disrupted by hA53T-α-Syn, while proteins including lysosomal proteins LAMP2 (lysosomal-associated membrane protein 2; increased) and GRN (progranulin; increased) MAPT (decreased), and immune proteins CD38 and FYN (both decreased) were impacted by the PFF-α-Syn model. Based on the representation of each disease-relevant process across *in vivo* models (Figure 10) and mechanisms of action for drug candidates (Figure 11A), we summarize recommendations for *in vivo* model selection (Figure 11B), wherein the AAV-hA53T model may be suited for testing therapeutics related to dopaminergic signaling or energy and mitochondria, which include GLP1R agonists, and the PFF model may be suited for testing therapeutics related to antioxidants, immune pathways, particularly glia-targeted therapeutics, or autophagy/endolysosome pathways, which include LRRK2-targeting therapeutics. In support of presence of neuroinflammation/immune function/ astrocyte biology in the PFF model, only the PFF-α-Syn model exhibited increases in the reactive astrocyte markers GFAP and S100B, and microglia marker ITGAM (integrin alpha M; CD11b) (Figure 7A right). Both models should be considered for testing neurotrophic factors or proteinopathy mechanisms, including α-Syn-or MAPT/Tau-targeted therapeutics.

**Figure 11:**
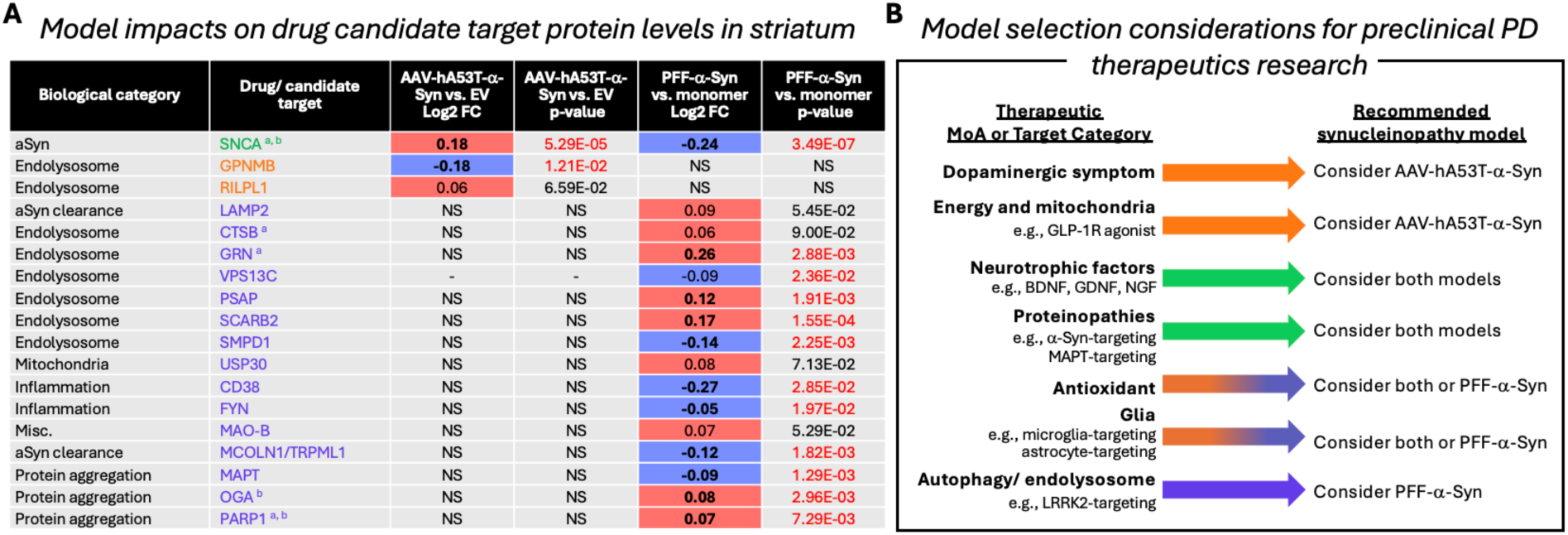
**Candidate therapeutic targets and considerations for PD drug development**. **A.** Proteins detected in striatum with changes in abundance (log2 fold change; p≤0.10) compared to respective control are shown; Those significantly altered (p≤0.05) appear in bold in either a red box (indicates increased abundance) or blue box (indicates decreased abundance). Significance of p≤0.05 is indicated by p-value in red. Protein names in green are impacted in both models, orange in hA53T-α-Syn, and purple in PFF-α-Syn. “-” indicates not detected; “NS” = not significant. Superscripts indicate significant (p≤0.05) differential abundance in human proteomics datasets, where a: PD vs control or b: DLB vs control (Shantaraman et al., 2024). **B.** A host of candidate therapeutic agents are currently in the development pipeline for the treatment of PD, many of which have mechanisms of action that may be relevant to study using the AAV-hA53T-and PFF-α-Syn rat models. The diagram summarizes the recommended preclinical model according to therapeutic mechanism, with examples of therapeutics within those categories, as supported by our multi-omics approach. Orange arrows indicate consideration of the AAV-hA53T-α-Syn model. Purple arrows indicate consideration of the PFF-α-Syn model, with the orange-to-purple gradient arrow indicating that while both models may be suitable, selection of the PFF model will be more appropriate. Green arrows indicate that both models should be considered.

## DISCUSSION

This is the first comparative comprehensive proteomic and transcriptomic characterization, to our knowledge, of the AAV-hA53T-α-Syn and PFF-α-Syn rat models of synucleinopathy. The results of our analysis bolster the hypothesis-driven analyses of these approaches that currently exist in the literature and provide a framework for comparison of these two models that may be useful to researchers seeking to probe a specific protein or mechanism of PD or dementia with Lewy bodies (DLB).

### Modeling α-Syn pathology in PD drug research

The advancement of disease-modifying therapeutics for PD has been challenged by the complex and multifactorial nature of the disease. There are currently well over 100 drug candidates in the clinical development pipeline for PD between Phases 1 and 3 – some aiming to address symptomology, while others aiming to impact a range of disease mechanisms to slow disease progression.^38^ Beyond symptomatic treatments, there are very few DLB-focused therapeutics under development that aim to address disease mechanisms, and include neflamapimod (NCT06815965; CervoMed) and zervimesine (CT1812; NCT05225415; Cognition Therapeutics^39^). Informed selection of animal models will improve efficiency and bolster the probability of success in demonstrate proof of mechanism or proof of concept with a given investigational therapeutic, as the animal model must successfully recapitulate the disease biology or mechanism under investigation the therapy intends to target or modulate. In PD, the accumulation of misfolded α-Syn leads to Lewy body formation and subsequent mitochondrial and lysosomal dysfunction, autophagy, oxidative stress, and neuroinflammation.^1^ These aberrant processes cause synaptic dysfunction and neuronal death, particularly dopaminergic neurons.^40^ Pathogenic α-Syn can also propagate between neurons, inducing these dysfunctional processes in neighboring cells, spreading the disease to previously healthy cells and tissue, contributing to the progressive nature of the disease.^41,42^ Given the significance of α-Syn in the pathological processes contributing to PD, and other synucleinopathies such DLB, modeling α-Syn in animals is an important step towards understanding the disease and developing therapeutics to disrupt these processes.

AAV-mediated delivery of mutant human A53T α-Syn and injection of PFF α-Syn are widely utilized approaches to introducing pathological α-Syn to rodent brain to model synucleinopathies and to recapitulate and evaluate the pathogenic synucleinopathy that occurs in PD. Overexpression of mutant human A53T α-Syn (by AAV-mediated delivery of transgene unilaterally), induces pathological accumulation of mutant A53T-α-Syn leading to nigral dopaminergic cell loss and motor deficits.^5,43^ As such, the AAV-A53T-α-Syn rat model is ideally suited to assess therapeutics investigating neuroprotective mechanisms of action in rescuing TH^+^ cell death and motor deficits.^44^ Injection of preformed fibrils (PFF-α-Syn) into the rat striatum to observe the propagation of α-Syn in PD leads to pathological “templating” resulting in reactive hyperphosphorylation of endogenous α-Syn and its progressive aggregation, a neuroinflammatory response including activation of microglia and reactive astrogliosis, and disruption in several homeostatic and quintessential physiological processes including autophagy, synaptic function and other neuronal functions.^7,8,45^ Thus, the PFF-α-Syn model is thought to be well-suited for the assessment of treatments aimed at interfering with spread of pathological α-Syn through the targeting of α-Syn release, uptake, and synaptic clearance, as well as other mechanisms. As these models have been used to successfully test therapeutics for PD (see introduction), they also may be useful for the broader class of synucleinopathies, including DLB, as these diseases exhibit overlap in pathology and in underlying biological dysfunction.^26^ A deeper understanding of the mechanistic impacts of each model may foster model selection to hopefully facilitate successful and expeditious drug discovery/development.

### Multi-omics approaches provide a unique perspective to bolster research and drug development

Pathway and comparative analyses of the disrupted proteomes in the α-Syn models validated previously discovered promising therapeutic targets and pathways disrupted in PD and other synucleinopathies, as well as illuminated novel proteins and pathways that these models represent to recapitulate key aspects of disease. Evaluation of proteins correlating with canonical PD-related outputs and disease hallmarks – loss of TH^+^ cell numbers, pS129-α-Syn pathology, and reduced dopamine levels – illuminated proteins most correlated to these key features of the disease. Further, comparison of the proteome of each model with those identified in human PD and DLB brain proteomic cohorts shed light on potential disease biomarkers^46^ and the translational value of these models.

### Proteomic and transcriptomic assessments of hA53T-and PFF-α-Syn rat brain highlight distinct and overlapping biological processes

Aspects of PD pathophysiology that drive key features of the disease, such as synapse and neurotransmission-related pathways, neural growth and remodeling, transport, protein localization, and stress response, were well-represented in the unbiased proteomic findings for each α-Syn model in agreement with the existing body of knowledge generated from hypothesis-driven studies.^4,7,47,48^ While some disease mechanisms were indeed prominent in both models, notably synapse and neurotransmission-related processes, other mechanisms were largely model-specific, such as secretory vesicle and autophagy pathways for the PFF-α-Syn model, and mitochondrial function and dopamine biology for the hA53T-α-Syn model (Figure 8). While dopamine was a predominant pathway in the unbiased AAV analysis but not the PFF analysis, interestingly, a clear dopamine signal was detected in the PFF when the commonly altered proteins were identified in the comparative analyses of proteins disrupted across both models (Figure 6). When dopamine correlates were identified for each model, a strong emphasis on vesicle and exocytosis-related processes in the PFF model was observed, whereas in the hA53T model, impacted proteins correlated with dopamine were more related to synapse organization and neural development (Figure 7) – highlighting subtle differences in dopamine-related pathways across models.

Synaptic processes were disrupted in both α-Syn models (Figure 10), in agreement with previous studies; for example, hA53T-α-Syn overexpression has been shown to reduce or redistribute expression of synaptic markers,^49^ and PFF-α-Syn is known to alter synaptic function and dendritic spine morphology prior to cell death.^50^ The unbiased proteomics approach facilitated the identification of new synapse-related proteins such as SV2C and NRXN2 (altered in both models), neuroligin 1 (NLGN1) (altered in hA53T-α-Syn) and neuronal pentraxin 1 (NPTX1), synaptotagmin 1 and 2 (SYT1 and 2), and the syntaxin family (altered in PFF-α-Syn). In previously reported network analyses of PD and DLB patient proteomes, the top significantly correlated pathways with disease included pre-and post-synaptic processes, including vesicular signaling and transport.^26^ Both *in vivo* models exhibit changes that recapitulate some key biological aspects of these diseases: synapse-related biological pathways were impacted by both approaches, but pathways specific to intracellular and synaptic vesicles and endocytosis/exocytosis/secretion were more prominent in PFF-α-Syn, and DA-related processes (dopaminergic synapses and biosynthesis) were more prominently impacted in the hA53T-α-Syn model. The more prominent impact of hA53T-α-Syn on dopamine neurons, and the accelerated timeline for cell death in this model, was also reflected in a greater impact on motor and muscle-related processes, in keeping with the canonical ability of mutant α-Syn overexpression to cause motor deficits^44^ and the known propensity of AAV-mediated hA53T-α-Syn overexpression to selectively impact dopaminergic cells.^5^

PFF-α-Syn proteomic changes showed enrichment in autophagy and vesicle-based secretion/ exocytosis pathways (Figure 10), which are pathogenic processes known to impact PD progression in patients: α-Syn accumulation leads to dysfunctional autophagy^51^ and vesicle secretion/ exocytosis is a key process by which pathogenic α-Syn spreads between neurons.^52^ Autophagy-related proteins and pathways were consistently impacted in the PFF-α-Syn proteome, bolstering the existing studies that have found that the α-Syn aggregation cascade initiated by PFFs leads to autophagy and subsequent neuronal dysfunction.^8,53^ Members of the syntaxin family, cathepsin family, and the Rab GTPase RAB33B – all implicated in PD^54–57^ – are central nodes in the prominent autophagy/ lysosome cluster in the PPI map of PFF-α-Syn impacted proteins in this study. Secretion and exocytosis-related themes were also prominent among the proteins impacted by PFF-α-Syn, a finding that is consistent with the underlying mechanism of the model, which involves secretion of α-Syn aggregates and propagation to neighboring cells.^7^ Furthermore, many of the secretion-related proteins found to be impacted by PFF-α-Syn are also synapse-related (syntaxin family members, SNAP29, and SLC18A2 (VAMP2)). Taken together, both *in vivo* models should be considered for drug development targeting synaptic signaling; however, the PFF model should be considered for therapeutic targets with roles in vesicle transport and endo/exocytosis/secretion, while the hA53T-α-Syn model should be considered for testing targets specific to dopaminergic synapses and dopaminergic neuroprotection.

Mitochondria and energy metabolism are also important processes that are impacted in PD pathophysiology.^58^ While we found energy metabolism to be somewhat impacted in the proteome of the PFF-α-Syn model, it was more prominently affected by hA53T-α-Syn, which also consistently showed an impact on mitochondria (Figure 10). This finding aligns with previous studies demonstrating that the hA53T mutant of α-Syn associates with mitochondria and induces mitochondrial dysfunction and cell death.^59–61^ In addition to providing additional evidence for this aspect of synucleinopathy in the hA53T-α-Syn model, our findings implicate specific proteins such as mitochondrial ribosomal proteins (MRPS family), mitochondrial fission process proteins (MTF family), and DAP3 (death-associated protein 3). These findings corroborate patient brain proteomics data, in which mitochondrial and ribosomal modules were observed as among the most strongly correlated with PD and DLB.^26^ Thus, given the human disease relevance, the hA53T *in vivo* model should be considered for pharmaceutical drug development targeting mitochondria or energy and metabolism.

Interpretation of these findings must consider the similarities and differences between the two models. The hA53T-α-Syn model induces overexpression of human mutant α-Syn whereas PFF-α-Syn introduces preformed fibrils of α-Syn to induce aggregation and propagation of endogenous α-Syn. Consistencies across models include same rat strain (Sprague Dawley), same sex (female), same age at introduction of pathological α-Syn, and same procedure (stereotaxic surgery), although region of injection differed. The proteins and pathways commonly disrupted across both models (compared to sham) might reflect pathways generally impacted in synucleinopathies, perhaps in both familial cases (where there is overexpression of mutant α-Syn) or in sporadic cases (where there is a lack of α-Syn increase) and may reflect a proteomic/ transcriptomic synucleinopathy signature. Furthermore, it is possible that proteins and pathways that differ between the models may reflect subtle distinctions between synucleinopathies (i.e., α-Syn overexpression vs. PFF templating) as well as heterogeneities across patients (such as substantia nigra vs. striatal etiology of pathogenesis, downstream brain regions impacted, and speed of progression to neurodegeneration.)

### Comparative analyses with human PD patient proteomes and PD drug development pipeline reveal disease-relevant biomarkers

Comparative analysis of the hA53T-or PFF-α-Syn rat proteomes against human PD and DLB patient proteomic datasets^26^ revealed a number of disease-relevant proteins altered in one or both of the α-Syn models that were altered in PD/ DLB patient brains. Of note, SV2C, GABA transporter 1 (SLC6A1), Nogo-66 receptor (RTN4R), and lysozyme 2 (LYZ2), found to be significantly impacted in the hA53T-α-Syn model as well as the human PD cohort, are central to disease-related processes as identified in the pathway analyses for hA53T-α-Syn (neurite outgrowth, immune function, neurodegeneration). Enrichment analysis of the overlap indicated disease-relevant pathways represented to be development/ growth factors and immune response, and overlapping proteins with the DLB patient cohort enriched for similar pathways related to immune response and neural development.

In the PFF-α-Syn model, neuronal pentraxin 1 (NPTX1), excitatory amino acid transporter 2 (SLC1A2), synaptotagmin 2 (SYT2), and SV2C expression levels were disrupted, as well as the human PD and DLB cohorts, and are central to disease-related processes impacted in PFF-α-Syn pathway analyses (synapses, synaptic vesicles, and neurotransmitter transport). Pathway analysis of proteins that were impacted in both the PFF-α-Syn model and in PD patients showed oxidative stress signaling and astrocyte biology in the top pathways, while the overlapping proteins with the DLB patient cohort confirmed that the PFF model captures the synaptic, neuronal remodeling, and apoptosis/ cell survival pathways thought to be altered in the disease. Interestingly, oxidative stress pathways were more prominent in the overlapping proteins between the PD cohort with the PFF-α-Syn model than with the hA53T-α-Syn model. Specifically, the pathways within oxidative stress identified and translationally relevant to PD involved mitochondrial SOD2, which is congruent with the observation of altered SOD2 levels in patients with synucleinopathies^62^ and risk of PD associated with SOD2 gene polymorphisms.^63,64^ These observations highlight that while our proteomics results indicate oxidative stress pathways altered in both hA53T-α-Syn and PFF-α-Syn models, the PFF model exhibits a specific facet of synucleinopathy pathophysiology. Additionally, the overlapping proteins between the PD cohort and PFF model were enriched in astrocyte biology as the top MetaCore pathway, which suggests that the PFF model recapitulates glial-mediated biology associated with PD that has been well-documented in PD and other synucleinopathies.^65^ Altered glial biology in the PFF model is also supported by the analysis of canonical inflammation markers, including elevated GFAP (Figure 7A), which is a widely used biomarker in clinical settings and trials for PD and DLB to assess neuroinflammation. MetaCore pathways found to be overlapping between the two *in vivo* models and across PD and DLB patient cohorts were related to Tau dysregulation, which may reflect the biological interaction between α-Syn and Tau regulation that has been observed in preclinical models,^66–69^ as well as the roles of both proteins in synaptic signaling, further underscored by the strong correlation of SNCA abundance and DLB network modules related to synaptic processes.^26^

Further investigation via comparative analyses of the *in vivo* model omics with patient datasets is warranted, given the differences in brain regions (striatal tissue for the *in vivo* models, versus frontal cortex tissue for the human cohorts). Additionally, there are differences in the total number of differentially abundant proteins significant for each model, which predispose datasets with more total significant proteins to have larger overlap with another dataset. For the AAV-hA53T-α-Syn model, overlaps with the PD and DLB patient brain proteomic analysis made up 11.9% and 18.8%, respectively, of the total significantly altered proteins (388). For the PFF-α-Syn model, overlaps with the PD and DLB patient brain proteomic analysis made up 12.4% and 19.8%, respectively, of the total significantly altered proteins (1550). Although proteins of interest are identified here through comparison of just two studies and require further investigation, validation of translational biomarkers has the potential to accelerate the early phases of drug development research.^46^

Finally, drug candidates already in development share mechanisms and targets that were reflected in the AAV-hA53T-or PFF-α-Syn proteomes (Figure 11A, B). Our analyses revealed an impact on specific proteins and transcripts that are targets of PD drug candidates: In addition to α-Syn itself, GPNMB and others significantly impacted by hA53T-α-Syn, and GRN, FYN, tau, and others significantly altered by PFF-α-Syn (Figure 11A). It is important to note that the individual protein changes (or lack thereof) in the animal models reflect abundance changes in the striatum, and protein changes may be region-specific and/or temporal. Pathway analyses and comparative analyses with patient proteomes extrapolated beyond individual protein targets to better inform recommendations for model selection (Figure 11B). For instance, while GLP1R was not an individual protein detected in the *in vivo* model or patient proteomics datasets, agents targeting GLP1R^38^ might be best suited for preclinical research in the hA53T-α-Syn model, which was found to impact mitochondria and energy metabolism consistently across our analyses and as precedented in preclinical proof of concept studies using this model.^23,70^ The proteomics analyses, taken together, support the selection of a particular model for a number of other therapeutic approaches. For example, there are 18 investigational therapeutics in the PD clinical development pipeline that target dopaminergic symptom relief.^38^ Preclinical studies of these and similar compounds may be better suited for the hA53T-α-Syn model, as DA-related proteins and processes were prominently impacted in this study. Additionally, compounds targeting neuroprotection may be better suited for the hA53T-α-Syn model given the rapid and robust loss of dopaminergic neurons.^23,70^ Conversely, agents targeting autophagy and/or endolysosomal pathways, including the LRRK2 agents currently in clinical trials, may have the best opportunity to demonstrate efficacy in the PFF-α-Syn model, and indeed have been previously been studied in this model (Figure 11B).^20,71,72^ The recommendations herein are a simplified starting point for model selection, and further research to elucidate each drug candidate’s mechanism of action must be taken into consideration for the appropriate model selection to test efficacy.

This is the first study to comprehensively evaluate the hA53T-and PFF-α-Syn rat models of synucleinopathy using multi-omics approaches, and replication of results in repeated studies will be needed to validate these findings. Further dissection of pathways involved in these *in vivo* models beyond the scope of our proteomics analyses will also be needed; for example, assessing enzyme activity and post-translational modification of proteins that dictate function. In sum, the results of this unbiased proteomics study of the AAV-hA53T-α-Syn and PFF-α-Syn models of PD synucleinopathy, along with the pathway, comparative, and correlation analyses of the impacted proteins have strengthened the body of knowledge of what key aspects of PD and DLB these models may capture and provide a starting point to assist with model selection based off the mechanism of interest. Comparative analyses with human PD patient proteomic data and the PD drug development pipeline revealed potential translational biomarkers disrupted preclinically and clinically, providing insights that may facilitate early-phase PD drug development and accelerate much-needed therapeutics for patients.

## Acknowledgements

We thank the members of the Atuka, Inc. scientific team, especially Drs. Tom Johnston and James Koprich for their expertise in the study design and execution of the in-life portions of these studies and model characterization-related analyses.

## Funding

This study was funded by the Michael J. Fox Foundation grant MJFF-021175 awarded to MEH (Cognition Therapeutics) in 2022.

## Data availability statement

The datasets generated during and/or analyzed during the current study are not publicly available but additional information may be provided by the corresponding author upon reasonable request.

## Notes

### Competing Interest Statement

BNL, AR, AOC, and MEH are employees and shareholders of Cognition Therapeutics. HAN, RS, and GL are consultants to Cognition Therapeutics. NTS, DD, and KP are co-founders, employees, consultants, and/or shareholders of EmTheraPro.

