## Supplemental materials for "Multi-omics comparative analyses of synucleinopathy models reveal distinct targets and relevance for drug development"

### SUPPLEMENTAL FIGURES

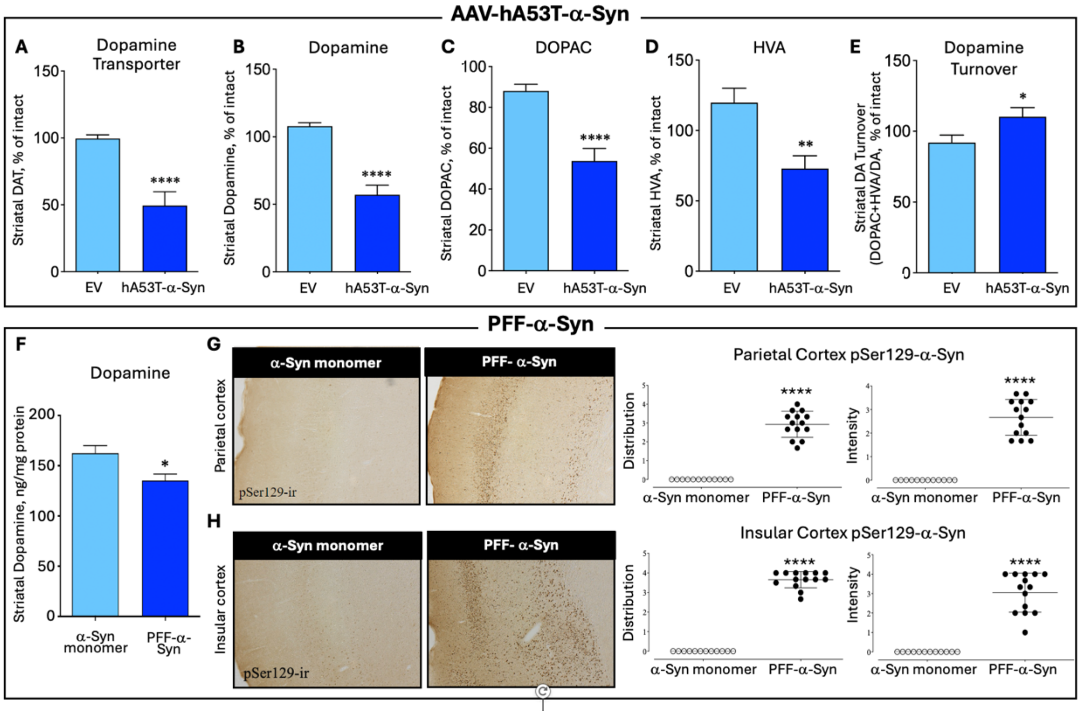


**Supplemental Figure 1. AAV-hA53T- and PFF-α-Syn models recapitulate key features of synucleinopathies. Top:** The impact of hA53T-α-Syn delivery was further confirmed through quantification (mean±SEM) of striatal dopamine transporter (DAT) **(A)**, dopamine **(B)**, DOPAC **(C)**, HVA **(D)**, and dopamine turnover (i.e., DOPAC+HVA/DA) **(E)**. **Bottom.** The impact of PFF-α-Syn delivery was further confirmed through quantification of striatal dopamine (mean±SEM) **(F)** and through parietal **(G)** and insular **(H)** cortex immunohistochemical (IHC) detection of α-Syn phosphorylated serine 129. Quantifications (mean±SD) of the distribution (percent of anatomical region of interest containing positive cells) and intensity (strength of IHC signal for each cell) of pSer129-α-Syn are shown.

**AAV-hA53T-α-Syn**

**A**

| GO Term | Biological Process | Strength | FDR |
|---|---|---|---|
| 0060997 | **Dendritic spine morphogenesis** | 0.97 | 2.28E-02 |
| 1900273 | **Positive regulation of long-term synaptic potentiation** | 0.88 | 4.86E-02 |
| 0016126 | Sterol biosynthetic process | 0.87 | 2.60E-03 |
| 1902653 | Secondary alcohol biosynthetic process | 0.87 | 4.40E-03 |
| 0006695 | Cholesterol biosynthetic process | 0.85 | 1.18E-02 |
| 0008045 | Motor neuron axon guidance | 0.84 | 1.34E-02 |
| 0097061 | Dendritic spine organization | 0.84 | 3.04E-02 |
| 2000273 | Positive regulation of signaling receptor activity | 0.82 | 3.58E-02 |
| 0060996 | Dendritic spine development | 0.81 | 3.90E-02 |
| 0017158 | Regulation of calcium ion-dependent exocytosis | 0.79 | 2.28E-02 |

Top STRING Biological Pathways (RNAseq) hA53T-α-Syn vs Empty Vector control (p≤0.05)

**B**

| Top Metacore Pathways (RNAseq) hA53T-α-Syn vs Empty Vector control (p≤0.05) | p-value |
|---|---|
| **G-protein signaling:** CDC42 inhibition | 9.22E-05 |
| **G-protein signaling:** Rap1A regulation pathway | 1.25E-04 |
| Signal transduction: Endothelin-1/ EDNRA signaling | 2.55E-04 |
| Activation of Ca2+-dependent neuronal cell death in Huntington's disease | 8.68E-04 |
| Development: Role of CDK5 in the nervous system | 1.69E-03 |
| Cell adhesion: Tight junctions | 1.71E-03 |
| Transport: Alpha-2 adrenergic receptor regulation of ion channels | 2.30E-03 |
| G-protein signaling: Rap2A regulation pathway | 4.27E-03 |
| Cholesterol Biosynthesis | 4.38E-03 |
| ATP metabolism | 4.38E-03 |

**C**

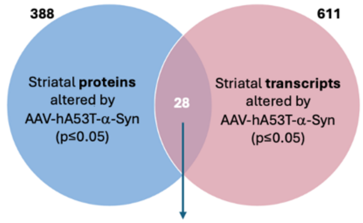


**D**

| Top Metacore Pathways (Overlap between Proteomics and RNAseq) hA53T-α-Syn vs Empty Vector control (p≤0.05) | p-value |
|---|---|
| **Axonal degeneration** in multiple sclerosis | 5.93E-04 |
| **Apoptosis and survival**: Regulation of apoptosis by mitochondrial proteins | 2.74E-03 |
| **Neurophysiological process: GABA-B receptor signaling at postsynaptic sides of synapses** | 1.95E-02 |
| Development: EGF-induced proliferation of Type C cells in SVZ of adult brain | 2.47E-02 |
| Activation of Ca2+-dependent neuronal cell death in Huntington's disease | 2.84E-02 |
| Sulfur metabolism | 3.13E-02 |
| Neurophysiological process: ACM2 and ACM4 signaling in the brain | 3.20E-02 |
| Neurophysiological process: Delta-type opioid receptor signaling in neuronal cells | 3.42E-02 |
| Regulation of lipid metabolism: PPAR regulation of lipid metabolism | 3.50E-02 |
| Neurophysiological process: Dopamine D2 receptor signaling in CNS | 3.50E-02 |

**PFF-α-Syn**

**E**

Top STRING Biological Pathways (RNAseq) PFF αSyn vs Monomer (p≤0.05)

| GO Term | Biological Process | Strength | FDR |
|---|---|---|---|
| 0045746 | Negative regulation of **Notch signaling pathway** | 0.92 | 6.60E-03 |
| 0035082 | Axoneme assembly | 0.9 | 1.26E-06 |
| 0006858 | Extracellular transport | 0.87 | 4.76E-02 |
| 0035850 | Epithelial cell differentiation involved in kidney development | 0.87 | 4.76E-02 |
| 0031670 | Cellular response to nutrient | 0.8 | 2.04E-02 |
| 0033628 | Regulation of cell adhesion mediated by integrin | 0.79 | 4.61E-02 |
| 0001578 | Microtubule bundle formation | 0.76 | 2.18E-05 |
| 0071277 | Cellular response to calcium ion | 0.67 | 2.70E-02 |
| 0003341 | Cilium movement | 0.66 | 4.00E-04 |
| 0060271 | Cilium assembly | 0.63 | 4.80E-07 |

**F**

| Top Metacore Pathways (RNAseq) PFF-αSyn vs Monomer (p≤0.05) | p-value |
|---|---|
| Development: **NOTCH signaling** inhibition | 1.02E-08 |
| Putative pathways of activation of **classical complement system** in major depressive disorder | 1.16E-06 |
| **Notch signaling** in oligodendrocyte precursor cell differentiation in multiple sclerosis | 1.50E-06 |
| Regulation **of immune cell differentiation** by **Notch signaling** | 5.23E-06 |
| Development: Inhibition of ossification by **NOTCH signaling** | 7.49E-06 |
| **Immune response: Classical complement pathway** | 1.12E-05 |
| Transcription: HIF-1 targets | 2.43E-05 |
| Development: **NOTCH signaling** activation | 4.45E-05 |
| **Immune response: Lectin induced complement pathway** | 6.64E-05 |
| Oxidative stress: ROS-induced cellular signaling | 7.38E-05 |

**Supplemental Figure 2: Transcriptomic analyses in A53T-α-Syn striatum and PFF-α-Syn substantia nigra. A-B.** Differential abundance of RNA transcripts, measured through RNA-seq detection in striatum, was assessed via ANOVA and significantly altered transcripts (log2 FC, AAV-hA53T-α-Syn vs. AAV-EV control; p≤0.05) were analyzed using STRING **(A)** and MetaCore **(B)** and show pathways affected in the AAV model. **C.** Comparative analysis of proteins (Figure 2) and transcripts altered in hA53T-α-Syn revealed 28 overlaps (i.e., protein and corresponding transcript). Of the 28 genes/proteins commonly altered, 19 were changing in the same direction (8 down, 11 up), whereas the remaining 9 were changed in opposing directions. **D.** MetaCore pathway analysis of these 28 overlaps revealed that apoptosis, neurodegenerative, and neuronal signaling processes were impacted in both the proteome and transcriptome of hA53T-α-Syn rats. **E-F.** Differential abundance of RNA transcripts, measured through RNA-seq detection in substantia nigra (SN), was assessed via ANOVA and 562 significantly altered transcripts (PFF-α-Syn vs. monomeric-α-Syn control; p≤0.05) were analyzed using STRING (v.11.5) **(E)** and MetaCore **(F)**. The significantly altered transcripts were enriched in pathways that include NOTCH, immune processes, and neural outgrowth/ remodeling. Processes of interest indicated in bold.

### SUPPLEMENTAL METHODS

*AAV-hA53T-α-Syn methods*

*Animals:* Thirty female Sprague-Dawley rats (Charles River, Quebec, Canada) weighing 248-319 g at the time of AAV surgery were used in this study. Animals were housed two per cage and acclimatized for at least one week prior to the start of any procedures. Animals were housed at standard temperature (21 ± 2$^{o}$C) in a light-controlled environment (lights on 6:00 a.m. to 6:00 p.m.) with access to food (Teklad 2020X, Harlan, Madison, WI) and water *ad libitum*. All protocols were approved by an independent Institutional Animal Care and Use Committee. Body weights were measured on D1 (prior to surgery), and weekly thereafter with the final measurement just prior to necropsy. Animals were assigned to experimental groups based on previously established methods:[21] the disease condition group receiving injection of AAV-hA53T-α-Syn to express human mutant α-Syn (referred to as AAV-hA53T-α-Syn hereafter) or the control group receiving injection of empty AAV vector (AAV-EV hereafter).

*AAV expression vector:* AAV1/2-hA53T-α-Syn utilizes the chimeric serotype 1/2. The A53T-α-Syn expressed by the gene delivered is of the human sequence. Its control was an empty AAV1/2 vector of the same serotype and viral construction. Both were diluted in sterile PBS and were administered at a volume of 2 µl; 0.5 µl/min. The concentration of AAV1/2 that was used (3 x $10^{12}$ genomic particles/ml) had previously been empirically determined to produce significant behavioral and dopaminergic nigrostriatal deficits between 3 and 6 weeks following surgical delivery (Koprich et al., 2011). AAV1/2 expression vector is referred to as AAV hereafter.

*AAV stereotaxic surgery:* Under isoflurane anesthesia (2% with 2 L/min oxygen flow rate, isoflurane USP 99.9%) the animal's head was shaved and the skin cleaned thoroughly using disinfectant soap, isopropyl alcohol 70% USP and iodine surgical scrub (7.5% povidone iodine). Rats were then placed in a Kopf small animal stereotaxic frame with the incisor bar set 3.3 mm below the ear bars (interaural line). An incision (~ 2 cm) was made with a sterile scalpel blade in an anteroposterior direction along the midline. After exposure of Bregma, a burr hole was drilled in the skull above the right SN at coordinates -5.2 mm AP, and -2 mm ML to Bregma (Paxinos and Watson, 1986). A customized 1 inch 26G Hamilton needle with 45-degree bevel was lowered -7.5 mm below the skull at the injection site. Animals received either AAV-hA53T-α-Syn or AAV-EV administered unilaterally into the right SN on Day 1 (D1). Following the injection, the needle was left in place for an additional 5 min to ensure complete absorption of the solution. After slow retraction of the injection needle, the incision was closed by means of wound clips and animals were administered saline (50 ml/kg, SC) and analgesic (Ketoprofen, 0.5 mg/kg, SC, ANAFEN®, Boehringer Ingelheim, Canada). The wound was coated with analgesic, antibacterial cream. Finally, animals were removed from the frame and placed in a recovery cage, positioned atop a thermostatically-controlled pad (HTP-1500 Heat Therapy Pump, Kent Scientific Corp., Torrington, CT, USA, set at 40ºC), and monitored until conscious.

*Behavioral assessment – cylinder test:* Cylinder test performance was assessed on D42 (6 weeks) relative to AAV administration. To examine animal forelimb asymmetry, rats were placed in a clear acrylic cylinder without top (15 cm diameter x 45 cm tall). To enhance rearing behavior, animals were deprived of food from 5 p.m. the previous night and testing was conducted between 8.30 a.m. and 11.30 a.m. the following morning. The number of times each paw touched the side of the cylinder during an individual rear was determined from *post hoc* analysis of video by an observer blinded to the treatments given. The first limb in any rear to touch the wall was scored a single point. If both limbs contacted within 0.4 s of each other, then this was scored as a 'both'. Following a single limb initially contacting the wall, all subsequent exploratory movements about the wall using that limb were scored independently until the other limb contacted the wall with weight support. Alternating stepping motions involving both paws one after the other received a single score for both. Asymmetry scores, %, were calculated using the formula: [100*(ipsilateral + both) - (contralateral + both) / (ipsilateral + both) + (contralateral + both)].

*Brain tissue processing:* Animals received an overdose of isoflurane and were sacrificed by exsanguination by transcardial perfusion with ice-cold 0.9 % saline containing 0.2 % heparin 24 ± 2 hours after the last daily dose. Brains were immediately removed and placed, ventral surface up, into an ice-cold stainless-steel brain matrix and first cut in the coronal plane at the level of the optic chiasma. A second cut was made 1 mm rostral to the first cut and striatum was freshly dissected for quantification of levels of DA and metabolites of DA (HVA and DOPAC) by liquid chromatography/mass spectrometry (LC-MS/MS), as described below. A third cut was made 1 mm rostral to the second cut and striatum was freshly dissected to assess the levels of human $\alpha$-Syn by ELISA, RNA sequencing, and proteomics, as described below. (Right (ipsilateral) hemisphere striatum was isolated for RNAseq and proteomics). The remaining rostral portion of the brain was immediately frozen in isopentane chilled to -42 °C, then stored at -80°C and later sectioned for DAT autoradiography. The caudal portion of the brain, including the mesencephalon, was immersed in 4% paraformaldehyde (PFA) for 48 hours for fixation, followed by cryoprotection in graded sucrose solutions (15 to 30 % sucrose). Tissue prepared in this manner was used for quantification of $TH^+$ neuron numbers in the SNpc via immunohistochemistry of tyrosine hydroxylase and unbiased stereology.

*Striatal DA and metabolites via LC-MS/MS:* Brain sections were homogenized, using a tissue dismembrator, in 100-750 µl of 0.1M TCA containing $10^{-2}$M sodium acetate, $10^{-4}$M EDTA, and 7.5% methanol (pH 3.8). 10 µl of homogenate was removed for measurement of protein concentration. The samples were then spun in a microcentrifuge at 10,000g for 20 minutes at 4°C. Supernatant was transferred to a new microcentrifuge tube for biogenic amine analysis. *Biogenic Amine Analysis:* DA, HVA and DOPAC were determined by a highly sensitive and specific LC-MS/MS methodology following derivatization of analytes with benzoyl chloride (BZC). 5 µl of supernatant was treated with 10 µl each of 500mM $NaCO_3$ (aq) and 2% BZC in acetonitrile. After four minutes, the reaction was stopped by the addition of 10 µl internal standard solution (in 20% acetonitrile containing 3% sulfuric acid) containing 200 pg of each $^{13}C_6$-derivatised dopamine-$d_4$, HVA, and DOPAC. LC was performed on a 2.0 x 50 mm, 1.7 $\mu$m particle Acquity BEH C18 column (Waters Corporation, Milford, MA, USA) using a Waters Acquity UPLC. Mobile phase A was 0.15% aqueous formic acid and mobile phase B was acetonitrile. Samples were separated by a gradient of 98–5% of mobile phase A over 11 minutes at a flow rate of 600 $\mu$l/min prior to delivery to a SCIEX 6500+ QTrap mass spectrometer (AB Sciex, Framingham, MA, USA). The following MRM transitions were monitored for quantitative purposes: 466 → 105, BZC-dopamine; 488 → 111, $^{13}C_6$-BZC-dopamine-$d_4$; 304 → 150, BZC-HVA; 310 → 111, $^{13}C_6$-BZC-HVA; 394 → 105, BZC-DOPAC; 406 → 111, $^{13}C_6$-BZC-DOPAC. Automated peak integration was performed using SCIEX MultiQuant software version 3.0.2. All peaks were visually inspected to ensure proper integration. Levels of DA, HVA and DOPAC in samples were calculated using calibration curves constructed on the basis of peak area ratio ($P_{analyte}/P_{I.S.}$) versus concentrations of internal standard by linear regression. Levels were normalized to protein concentration in the tissue extract. Protein concentration in tissue homogenates was determined using the Pierce™ BCA Protein Assay Kit (Thermo Fisher Scientific, Waltham, MA USA) as described in the provided kit instructions. Absorbance was measured using a POLARstar Omega plate reader (BMG LABTECH, Offenburg, Germany).

*$\alpha$-Syn overexpression:* Freshly dissected striatal tissue of all animals was homogenized in a lysis buffer containing protease and phosphatase inhibitors (Roche: 11836153001). Samples were agitated at 4°C for 30 minutes followed by centrifugation (135000 rpm for 10 minutes at 4°C) to produce supernatant. Using a 1:500 dilution for a concentration of 0.001 mg/ml, a portion of supernatant was used to determine total protein levels (BCA assay, Pierce, Rockford, IL). The remaining supernatant underwent ELISA procedures according to the manufacturer's instructions (BioLegend: 844101). The samples were analyzed using CLARIOstar systems quantifying the luminescent counts relative to the amount of $\alpha$-Syn. Levels of $\alpha$-Syn

were expressed as pg/mg total protein (PierceTM BCA Protein Assay Kit, Thermo Fisher Scientific, Waltham, MA USA).

*DA transporter (DAT) binding:* The levels of striatal DAT were assessed in all animals by [$^{125}$I]-RTI-121 binding autoradiography in cryostat cut sections prepared from 20 µm fresh-frozen tissue. Briefly, thawed slides were placed in binding buffer (50 mM Tris, 120 mM NaCl and 5 mM KCl, 2 x 15 minutes, room temperature). Sections were then placed in the same buffer containing 1 pM [$^{125}$I]-RTI-121 (Perkin-Elmer, specific activity 550 Ci/µmol) for 90 minutes at room temperature to determine total binding. All slides were then washed (2 x 20 minutes) in ice-cold PBS buffer, rinsed in ice-cold distilled water and air-dried. Together with [$^{14}$C] microscale standards (Amersham) slides were then apposed to autoradiographic film (Kodak) and left for 4 days at room temperature before developing. Autoradiograms were then analyzed using MCID software (Image Research Inc, Ontario, Canada). Densitometric analysis of 3 striatum cryosections from each animal was carried out whereby a reference curve of c.p.m. versus optical density was calculated from β-emitting [$^{14}$C] microscale standards and used to quantify the intensity of signal as nCi/g. Background intensity was subtracted from each reading. Data were then expressed as mean ± SEM signal intensity for each treatment group.

*Histology:* A single series of sections from the midbrain were processed for visualization of tyrosine hydroxylase (TH) via the biotin-labelled antibody procedure to label DA neurons of the SN. Briefly, following several washes in a PBS solution containing 0.2% Triton X-100 (PBS-T), endogenous peroxidase was quenched in a 3% hydrogen peroxide solution and background staining inhibited in a 10% normal goat serum/2% bovine serum albumin solution. Tissues were then incubated with primary antibodies overnight: rabbit anti-TH antibody (1:1000, Millipore, AB152). After three washes in PBS-T, sections were sequentially incubated in biotinylated goat anti-rabbit or mouse IgG (1:400; Jackson Immuno, 111-065-144) for 1 h and the Elite avidin-biotin complex (ABC Kits; Vector, Burlingame, CA) for 1 h separated by three washes in PBS. Immunostaining was then visualized following a reaction with 3,3-diaminobenzidine (Vector, Burlingame, CA). Sections were mounted on glass slides, allowed to dry, dipped into dH20, dehydrated through graded alcohols (70%, 95%, 100%), cleared in xylenes, and coverslipped with DPX mounting medium (Electron Microscopy Sciences, Hatfield, PA).

*Stereology:* Estimates of $TH^+$ neuronal number within the SNpc were performed using Stereo Investigator software (MBF Bioscience, Williston, VT) according to stereological principles. Eight sections, each separated by 140 µm from the anterior to the posterior SN, were used for counting each case. Stereology was performed using a Zeiss microscope (Carl Zeiss, Canada) coupled to a digital camera for visualization of tissue sections. The total number of $TH^+$ neurons was estimated from coded slides using the optical fractionator method. For each tissue section analyzed, section thickness was assessed empirically and guard zones of ~2 µm thickness were used at the top and bottom of each section. The SNc was outlined under low magnification (5x) and $TH^+$ neurons counted under 40x magnification. Stereological parameters were empirically determined (i.e. grid size, counting frame size and dissector height) using Stereo Investigator software (MicroBrightfield, VT, USA). The acceptable coefficient of error (CE) was calculated according to the Gunderson CE (m=1) procedure. Gunderson values < 0.10 were accepted.

*Statistical analyses:* Continuous data derived from calculations of forelimb asymmetry and all *postmortem* endpoints (including striatal DA and metabolite levels, DAT, human transgene-derived $\alpha$-Syn levels, and nigral $TH^+$ cell counts) were graphed as mean ± SEM. Group comparisons between AAV-EV and AAV-hA53T-α-syn animals were performed using a two-tailed student's unpaired t-test. $TH^+$ cell counts were subjected to correlational analyses with the forelimb asymmetry test at week 6, DA levels and its metabolites for both treatment groups.

*PFF-$\alpha$-Syn study methods*

*Animals* Twenty-six female Sprague-Dawley rats (Charles River, Quebec, Canada) weighing 251-319 g at the time of PFF surgery were used in this study. Animals were housed, fed, and acclimatized prior to procedures, and handled according to protocols approved by IACUC as described above for AAV surgeries. Animals were assigned to experimental groups: the disease condition group receiving injection of mouse $\alpha$-Syn PFFs or the control group receiving mouse monomeric $\alpha$-Syn.

*Surgery* Animals received either mouse PFF-$\alpha$-Syn (mPFF Type-1, StressMarq, 8 µg/ml) or monomeric $\alpha$-Syn control administered bilaterally into the striatum according to stereotaxic techniques on D1 (2 x 2 µl of 8 µg/µl mPFF = 32 µg total). The injection sites were +1.6 mm AP and +/- 2 mm ML with the needle lowered 4 mm from skull (site 1) and -0.1 mm AP, +/- 4.2 mm ML with the needle lowered 5 mm from skull (site 2). (2 x 2 µl of 8 µg/µl mPFF = 32 µg total). The injection sites were +1.6 mm AP and +/- 2 mm ML with the needle lowered 4 mm from skull (site 1) and -0.1 mm AP, +/- 4.2 mm ML with the needle lowered 5 mm from skull (site 2). All surgeries were performed using aseptic technique. Under isoflurane anesthesia (2% with 2 L/min oxygen flow rate, isoflurane USP 99.9%) and after confirmation of loss of tail-pinch and corneal reflexes, rats were placed in a Kopf small animal stereotaxic frame with the incisor bar set 3.3 mm below the ear bars (interaural line). The animal's head was shaved and the skin cleaned thoroughly using disinfectant soap, isopropyl alcohol 70% USP and iodine surgical scrub (7.5% povidone iodine). An incision (~ 2 cm) was made with a sterile scalpel blade in an anteroposterior direction along the midline. After exposure of Bregma using a cotton-bud, a burr hole was drilled in the skull above the striatum (Day 1) at the coordinates described above according to the atlas of Paxinos and Watson (1986). A customized, 1 inch, 26G Hamilton needle with 45-degree bevel will be used for all injections. After each injection the needle was left in place for an additional 5 min to ensure complete absorption of the solution. After slow retraction of the injection needle, the incision was closed by means of wound clips and animals were administered saline (50 ml/kg, SC) and analgesic (Ketoprofen, 0.5 mg/kg, SC, ANAFEN®, Boehringer Ingelheim, Canada). The wound was coated with analgesic, antibacterial cream. Finally, animals were removed from the frame and placed in a recovery cage, positioned atop a thermostatically-controlled pad (HTP-1500 Heat Therapy Pump, Kent Scientific Corp., Torrington, CT, USA, set at 40oC), and monitored until conscious.

*Brain tissue processing:* On D60, animals were deeply anaesthetised with isoflurane and killed via exsanguination by way of transcardial perfusion with ice-cold 0.9 % saline containing 0.2 % heparin. Brains were then placed, ventral up, into an ice-cold stainless steel rat brain matrix and first cut was in the sagittal plane bisecting the right and left hemispheres. A second cut was made at the level of the optic chiasm on the left side of the brain. The remaining right hemisphere was immersed in 4% paraformaldehyde (PFA) for 48 hours fixation, followed by cryoprotection in graded sucrose solutions (15 to 30 % sucrose) for immunohistochemistry (pSer129-aSyn). The left forebrain was flash frozen in isopentane chilled to -42 °C. Striatal tissue was later dissected from cryosections for quantification of DA and its metabolites via LC/MS analysis (2 x 80 µm sections), DAT by autoradiography (3 x 20 µm sections; optional) and remaining tissue for proteomics (remaining x 80 µm sections). Left hemisphere SN was harvested for RNAseq. The remaining caudal portion of the left hemisphere was flash frozen in isopentane chilled to -42 °C and later ventral midbrain was dissected from cryosections for RNAseq.

*pSer129 $\alpha$-Syn immunohistochemistry:* Post fixed and cryo-preserved sections of forebrain and midbrain were processed for pSer129 $\alpha$-Syn immunoreactivity. Brains were cut frozen in the coronal plane at a thickness of 40 µm on a sledge microtome (Leica) and 6 series of sections were stored in cryoprotectant. One series of sections was processed for visualization of pSer129 $\alpha$-Syn via the biotin-labeled antibody procedure. Briefly, following several washes in a PBS solution containing 0.1% Tween-20, endogenous peroxidase was quenched in a 3% hydrogen peroxide solution. Antigen retrieval was performed by incubating the sections in a sodium citrate buffer for 1 hr at 37o C followed by 3 rinses in PBS. Background staining was inhibited in a 5% normal donkey serum / 2% bovine serum albumin solution. Tissue was then incubated overnight with anti pSer129 $\alpha$-Syn (1:5000; Abcam, ab51253). After three washes in PBS, sections were sequentially incubated in biotinylated antibodies IgG (donkey anti-rabbit, 1:500; Jackson, West Grove, PA) and the Elite avidin-biotin complex (ABC Kits; Vector, Burlingame, CA) for 1 h separated by three washes in PBS. pSer129 $\alpha$-Syn immunostaining was visualized following a reaction using 3,3-diaminobenzidine (ABC Elite, Vector Laboratories, Cat. # PK-6100). Sections were then mounted on glass slides, allowed to dry, dipped into dH20, dehydrated through graded alcohols (70%, 95%, 100%) and cleared in histo-clear prior to being coverslipped with Vecta-mount mounting medium. Following intrastriatal injection in rats, pathological $\alpha$-Syn formation and levels have been shown to peak in many cortical regions within 1-2 months, with nigrostriatal degeneration beginning at 2-4 months.[12,13]

*Cell counts:* Estimates of pSer129-$\alpha$-Syn$^+$ cell numbers within the SNpc were performed using Stereo Investigator software (MBF Bioscience, Williston, VT) by technicians blinded to treatment. Three sections, equally spaced from the anterior to the posterior SN, were used for counting each case. All cells observed were counted using a Zeiss microscope (Carl Zeiss, Canada) coupled to a digital camera for visualization of tissue sections. The total number of cell profiles were taken from 5x magnification. The average of the counts from 3 sections are reported±SEM.

*Qualitative assessment of pSer129 $\alpha$-Syn signal in the forebrain:* The initial analyses of pSer129-$\alpha$-Syn$^+$ cell signal intensity and distribution were qualitatively assessed using a Zeiss microscope (Carl Zeiss, Canada) in the parietal and insular cortices of rat forebrain sections at 3 anterior-posterior brain levels relative to Bregma: 1.6 mm, -0.6 mm, and -1 mm. The intensity of pSer129-$\alpha$-Syn$^+$ cells was rated based on the following criteria: 0 = no signal; 1 = faint pSer129-$\alpha$-Syn$^+$ signal; 2 = 2x stronger pSer129-$\alpha$-Syn$^+$ signal than a 1 score; 3 = 3x stronger pSer129-$\alpha$-Syn$^+$ signal than a 1 score; 4 = 4x stronger pSer129$^+$ signal than a 1 score. The distribution of pSer129-$\alpha$-Syn$^+$ cells was rated based on the following criteria: 0 = no signal; <1 = pSer129-$\alpha$-Syn$^+$ signal in less than 25% of the region of interest (ROI); 1 = pSer129-$\alpha$-Syn$^+$ signal in 25% of the ROI; 2 = pSer129-$\alpha$-Syn$^+$ signal in 50% of the ROI; 3 = pSer129-$\alpha$-Syn$^+$ signal in 75% of the ROI; 4 = pSer129-$\alpha$-Syn$^+$ signal in 100% of the ROI. The final scores were averaged over 3 sections by a technician blinded to treatment; each dot on the plot (Figure 1B and Supplemental Figure 1G, H) represents the average score of the 3 sections per animal from the target region within the tissue section. The insular cortex and parietal cortex have previously been shown to harbor a rich density of pSer129 immunoreactivity in model characterization studies conducted in house. Figure 1B and Supplemental Figure 1G-H show representative data using the qualitative method of p129 assessment. Graphs are of mean±SD, one-way ANOVA with uncorrected Fisher's post hoc test.

*Brain Tandem-mass tag mass spectrometry (TMT-MS) proteomics measurements*

Brain samples were processed and analyzed using Tandem-mass tag mass spectrometry (TMT-MS) followed by unbiased quantification (AAV study: N = 18 animals, 10 = empty vector (AAV-EV), 8 = AAV-hA53T-$\alpha$-Syn; PFF study: N = 20 animals, 10 = monomer, 10 = PFF-$\alpha$-Syn).

*High-pH peptide fractionation*: High-pH peptide fractionation was conducted as previously described (Lizama et al., Neurobiol Dis. 2024). Dried samples were re-suspended in high pH loading buffer (0.07% vol/vol NH4OH, 0.045% vol/vol FA, 2% vol/vol I) and loaded onto a Water's BEH column (2.1 mm × 150 mm with 1.7 µm particles). A Vanquish UPLC system (Thermo Fisher Scientific) was used to carry out the fractionation. Solvent A consisted of 0.0175% (vol/vol) NH4OH, 0.01125% (vol/vol) FA, and 2% (vol/vol) I; solvent B consisted of 0.0175% (vol/ vol) NH4OH, 0.01125% (vol/vol) FA, and 90% (vol/vol) I. The sample elution was performed over a 25 min gradient with a flow rate of 0.6 ml/min with a gradient from 0 to 50% solvent B. A total of 192 individual equal volume fractions were collected across the gradient. Fractions were concatenated to 96 fractions and dried to completeness using vacuum centrifugation.

*Mass spectrometry analysis and data acquisition:* All samples (~1 µg for each fraction) were loaded and eluted by an Ultimate U3000 RSLCnano (Thermo Fischer Scientific) with an in-house packed 15 cm, 150 µm i.d. capillary column with 1.7 µm CSH (Water's) over a 35 min gradient. Mass spectrometry was performed Orbitrap QE-HFX (Thermo Fisher) in positive ion mode using data-dependent acquisition with Top 20 cycles. Each cycle consisted of one full MS scan followed by as many as 20 MS/MS. MS scans were collected at a resolution of 120,000 (410–1600 m/z range, 3×10^6 AGC, and 50 ms maximum ion injection time). Only precursors with charge states between 2+and 5+ were selected for MS/MS. All higher energy collision-induced dissociation (HCD) MS/MS spectra were acquired at a resolution of 45,000 (0.7 m/z isolation width, 32% collision energy, 1×10^5 AGC target, 120 ms maximum ion time). Dynamic exclusion was set to exclude previously sequenced peaks for 10 s within a 10-ppm isolation window.

*Database search and protein quantification:* All raw files were analyzed using the Proteome Discoverer Suite (v.2.4.1.15, Thermo Fisher). MS/MS spectra were searched against the UniProtKB rat proteome database (downloaded in May 2022 with 29918 total sequences). The Sequest HT search engine was used to search the RAW files, with search parameters specified as follows: fully tryptic specificity, maximum of two missed cleavages, minimum peptide length of six, fixed modifications for TMTPro tags on lysine residues and peptide N-termini (+304. 304.2071 Da) and carbamidomethylation of cysteine residues (+57.02146 Da), variable modifications for oxidation of methionine residues (+15.99492 Da), histidine, serine, threonine and tyrosine TMTPro tags (+304.2071 Da) and deamidation of asparagine and glutamine (+0.984 Da), precursor mass tolerance of 10 ppm and a fragment mass tolerance of 0.05 Da. Percolator was used to filter peptide spectral matches and peptides to an FDR<1%. Following spectral assignment, peptides were assembled into proteins and were further filtered based on the combined probabilities of their constituent peptides to a final FDR of 1%. Peptides were grouped into proteins following strict parsimony principle.

*Total RNA extraction and RNA sequencing*

Total RNA was extracted from striatal tissue (AAV model) and substantia nigra (PFF model) using the RNeasy kit (Qiagen, Germany) following the manufacturer's protocol ((AAV study: N = 18 animals, 10 = empty vector (EV), 8 = AAV-hA53T-$\alpha$-Syn; PFF study: N = 18 animals, 10 = monomer, 8 = PFF-$\alpha$-Syn). RNA quality was assessed using the Agilent Bioanalyser, quantified using Qubit (Thermo Fisher Scientific) and high-quality samples (RIN > 8) selected for sequencing analysis. RNA-seq library preparation, sequencing and bioinformatic analyses including differential expression analysis was performed at Azenta Life Science (South Plainfield, NJ, USA). Using DESeq2, a comparison of gene expression between AAV-hA53T-$\alpha$-Syn vs AAV-EV and between PFF-$\alpha$-Syn vs monomer-$\alpha$-Syn was performed. The Wald test was used to generate p-values and log2 fold changes. A total of 14,879 transcripts were detected reliably across AAV model samples, and 14,932 transcripts were detected reliably across the PFF model samples.

*Pathway analyses of significantly impacted proteins and genes*

<u>GO terms categorization analysis:</u> The Biological Process GO terms were categorized as relating to synapse and neurotransmission, dopaminergic synapses or DA biosynthesis, neuronal growth and remodeling, vesicles and secretion, protein degradation, autophagy, or others, and quantified accordingly. The Cellular Component GO terms were categorized as relating to either synapses and neurotransmission, dopaminergic synapses, neuronal growth and remodeling, vesicles and secretion, protein degradation, autophagosome, or mitochondria, and quantified accordingly. Reactome terms were categorized as relating to either synapses and neurotransmission, neuronal growth and remodeling, vesicles and secretion, protein degradation, immune system, and mitochondria or metabolism, and quantified accordingly.
